# Global geography outweighs long-term dynamics in shaping marine microbial population structure

**DOI:** 10.64898/2025.12.20.695618

**Authors:** Francisco Latorre, Lidia Montiel, Vanessa Balagué, Ramon Massana, Josep M. Gasol, Pierre E. Galand, Ramiro Logares

## Abstract

Marine microbial populations play essential roles in ocean ecosystems, yet the processes shaping their genomic structure across space and time remain poorly understood. Here, we examined the population-scale patterns of 1,505 prokaryotic metagenome-assembled genomes (MAGs) retrieved from the Northwestern Mediterranean Sea in two long-term coastal time series, along 15 and 7 years in these sites, as well as in the global ocean. We found that populations were generally more genomically differentiated across large spatial scales than across long temporal scales. Among a subset of 389 MAGs well represented in all datasets, 68.4% showed weak population divergence over time but strong global-scale differentiation. This was evident in the abundant and widespread cyanobacteria *Prochlorococcus* and *Synechococcus*. In contrast, only 6.4% of the MAGs exhibited weak divergence over time and space, with SAR11 MAGs being a clear example, likely reflecting their high dispersal and recombination rates. Other groups, such as SAR86 and Flavobacteriales, showed strong divergence at temporal and spatial scales, suggesting seasonal and/or regional adaptation. Positive selection was more readily detectable in the long-term coastal observatories than in the global ocean, despite the more significant population divergence observed across broad geographic scales. Temperature consistently showed a significant association with the population structure of many MAGs. Overall, our results highlight the dominant influence of large geographic scales in shaping microbial population structure alongside taxon-specific responses to temporal variation, particularly seasonality. Altogether, our work advances the understanding of microbial population structure across broad spatial and temporal scales, a critical step toward predicting microbial dynamics in a changing ocean.

## INTRODUCTION

Accounting for approximately 10^30^ cells^1^, prokaryotes are among the main contributors to the global microbial biomass^2^. In the ocean, bacteria and archaea play key roles in the biogeochemical cycling of nutrients, matter, and energy^3,4^. Marine prokaryotes are highly diverse and encompass various lineages that are able to perform a wide array of complex metabolic processes^5^, allowing them to colonize different habitats, such as pelagic zones, subsurface open-ocean waters, and sediments^6^. Despite their importance, key aspects of the diversity, ecology, and evolution of marine prokaryotes remain poorly known^7^. In particular, our understanding of prokaryotic populations, defined as individuals belonging to the same species, remains an underexplored dimension of biodiversity. Overall, we still lack a solid understanding of the diversity and structure of wild microbial populations and the ecological and evolutionary processes that shape them^8^.

Advancing our understanding of microbial population diversity, structure, and adaptations is crucial for elucidating the function of marine ecosystems and for improving predictions of microbiome responses to global change. Fundamental questions that remain partially answered are: How much genetic diversity is present within marine microbial species? What are the main drivers shaping their population structure? And how do populations adapt to environmental heterogeneity (*i.e.,* what genomic mechanisms underlie population adaptation across species?).

Over the past decades, it has become increasingly clear that the major drivers shaping marine microbial communities include oceanographic features, such as depth, currents, and water masses, along with the physicochemical properties of seawater^9–12^. In contrast, the factors shaping microbial population structure remain largely unknown. This is accentuated by the ongoing debate over what constitutes a “species” in prokaryotes, since neither phenotypic nor genomic traits provide clear boundaries between populations and species^13,14^. Oceanographic features, including water masses and currents, can limit or promote microbial dispersal between oceanic regions or depths^15^, where physical and chemical variables such as temperature, salinity, and nutrient availability may differ significantly. These environmental differences can exert heterogeneous selection, which, coupled with varying levels of dispersal, may drive the adaptation of microbial populations to distinct conditions found in specific basins or at different depths^16,17^. The large population sizes and relatively rapid reproductive rates of microbes are expected to facilitate their adaptation to local or regional environmental conditions, compared with animals and plants^18^. Such adaptations could happen in contemporary or ecological timescales^19^. Yet, marine microbes may also display rapid adaptation in constant conditions^20,21^ and may evolve faster through non-adaptive (neutral) processes over large spatial scales^22^.

Ideally, microbial population genomics would investigate genomes from multiple individuals or cells. However, obtaining full genomes of environmental microbes is challenging, as most remain uncultured^6,23^. Over the past decade, metagenomics as well as single-cell genomics have allowed the retrieval of partial or full genomes from uncultured microorganisms^24–29^. These genomes, known as Metagenome-Assembled Genomes (MAGs) or Single-Amplified Genomes (SAGs), typically have lower quality than gold-standard genomes obtained from cultures^30,31^. Yet, they provide access to the genomic information of microbes that would otherwise remain inaccessible. These genomes can subsequently be used in comparative population genomics or integrated with metagenomes to reconstruct population-level variation^8,32^.

Despite the increasing availability of marine MAGs, SAGs, and metagenomes^27,29,33,34^, relatively few studies have utilized them to investigate the structure of microbial populations and their potential adaptations to local or regional environmental conditions. Among the few available studies, those involving the SAR11 group, *Prochlorococcus*, and *Synechococcus* stand out, showing that population structure has not been influenced by dispersal limitation but instead correlates with temperature, nutrient, and light availability over different ocean regions^28,35–38^. Some SAR11 populations exhibit limited changes in abundance across marine niches, a pattern underpinned by high rates of homologous recombination between closely and distantly related lineages. This extensive genetic exchange promotes population homogenization, resulting in a stable population that may remain unchanged, yet exhibits high intrapopulation diversity over extended temporal and spatial scales^39^. Genomic differentiation among specific SAR116 populations has also been reported, with some considered endemic to the Mediterranean Sea and believed to be adapted to particular environmental conditions, such as low phosphate concentration^40^. Other investigations on hydrothermal vents have revealed significant genomic variation within microbial populations, indicating distinct evolutionary pressures in different geochemical settings^41^.

To date, most studies of marine microbial populations have treated spatial and temporal patterns as separate entities. Here, we examine both dimensions together to advance understanding across scales. For that, we investigate the population structure of 1,505 marine MAGs, each representing a potential species in two unique long-term time series spanning 15 and 7 years, as well as in the surface global ocean. These MAGs were recovered from the Long Term Ecological Research (LTER) Blanes Bay Microbial Observatory (BBMO)^42^, a coastal site in the Northwestern Mediterranean Sea. We tested whether MAG population diversity, structure, and potential adaptations vary across spatiotemporal scales, and examined the role of environmental factors in shaping populations. To do so, we first compared the monthly population variation of the selected MAGs at two geographically close long time-series sites: the LTER BBMO, during 15 years, and the Banyuls Bay Microbial Observatory (SOLA)^43–45^, spanning 7 years. We then investigated spatial population patterns of the 1,505 MAGs in the global ocean using a broad metagenome dataset from the *Tara Oceans* expedition (2009–2013)^46–48^.

## METHODS

### Sampling, DNA extraction, and shotgun sequencing

Coastal surface water samples (1 m depth) were collected at the Blanes Bay Microbial Observatory (BBMO, 41°40′N, 2°48′E; http://bbmo.icm.csic.es) in the Bay of Blanes, Spain. BBMO is an oligotrophic coastal site located about 1 kilometer (km) offshore, with a depth of approximately 20 meters and minimal influence from rivers or human activity^42^. Furthermore, samples were also collected at the Banyuls Bay microbial observatory (SOLA, 42°29’N, 3°08’E) in the Bay of Banyuls-sur-Mer, France. SOLA is also an oligotrophic coastal site, situated approximately 1 km off the coast of Banyuls-sur-Mer, France, with a depth of around 26 meters, and minimal human impact^49^. However, unlike BBMO, SOLA experiences sporadic winter storms, which resuspend nutrients from the sediments into the water column. Additionally, flash floods from nearby rivers further enrich SOLA’s waters with nutrients^43^. Both stations are separated by ∼150 km in the Northwestern Mediterranean Sea and connected by a dominant south-western marine current, also known as the Liguro-Provençal-Catalan Current. This current travels along the Italian coast west of Genoa, as well as the French and Catalan coastlines, with surface speeds that can reach 1 m/s^50^. Despite being a stable current, in the summer months it loses its strength to the benefit of coastal water masses flowing from south to north of the Iberian Peninsula, potentially increasing water-mass differentiation between BBMO and SOLA^51^.

In BBMO samples, ∼5 L of 200-μm pre-filtered surface seawater were sequentially filtered through a 20-μm mesh, a 3-μm pore-size polycarbonate filter (Poretics), and a 0.22-μm Sterivex Millipore (Merck-Millipore) filter using a peristaltic pump. At SOLA, a subsample of 5 L from 10 L Niskin bottles was sequentially filtered through 3 μm pore-size polycarbonate filters (Merck-Millipore, Darmstadt, Germany) and 0.22-μm Sterivex Millipore (Merck-Millipore) filters. Sterivex cartridges containing the pico-fraction (0.22–3 μm) of the microbial biomass were stored at –80 °C until nucleic acid extraction for both sampling stations. Note that sampling at both locations was part of the regular sampling at each time series. Monthly samples from January 2008 to December 2022 for BBMO and from January 2009 to December 2015 for SOLA (15 and 7 years, respectively) were used, yielding a dataset of 175 samples for the BBMO station and 90 samples for the SOLA station (**Supplementary Dataset S1**).

DNA extractions of the BBMO samples were performed according to the protocol described by Schauer *et al.*^52^, in which a lysozyme solution (20 mg/mL) was added to the Sterivex cartridges to lyse cells. Subsequently, a second incubation with proteinase K (20 mg/mL) was applied. The pooled lysates were then extracted twice with an equal volume of phenol-chloroform-isoamyl alcohol (25:24:1, pH 8), and once with an equal volume of chloroform-isoamyl alcohol (24:1). Finally, a concentration and purification step were performed using Amicon 100 (Millipore) units. DNA extractions of the SOLA samples were performed according to the protocol described by Hugoni *et al.*^53^, using a lysozyme solution (20 mg/mL) for the cell lysis with a second incubation with proteinase K (20 mg/mL), and the pooled lysates were then extracted using the AllPrep DNA/RNA kit (Qiagen, Hilden, Germany).

Before sequencing the BBMO metagenomes, DNA quality control was performed using an agarose gel and a NanoDrop One Spectrophotometer (Thermo Scientific). DNA quantification was performed using a Qubit fluorometer. From January 2009 to December 2011 (3 years), samples were sequenced on an Illumina HiSeq 4000 (2 x 150 bp) platform, and from January to December 2008 (1 year), and January 2012 to December 2022 (11 years) on an Illumina NovaSeq 6000 (2 x 150 bp) at the Centre Nacional d’Anàlisi Genòmica CNAG, Barcelona, Spain. About 45 billion reads were produced. For sequencing SOLA metagenomes, DNA quality control was performed using the Agilent High Sensitivity kit (Agilent Technologies, Santa Clara, CA, USA). Between January 2009 and December 2011 (3 years), the samples were sequenced on an Illumina NovaSeq 6000 platform (2 × 150 bp) at the Centre Nacional d’Anàlisi Genòmica (CNAG), Barcelona, Spain. Between January 2012 and February 2015 (∼3 years), the samples were sequenced on an Illumina HiSeq 2500 platform (2 × 100 bp) at GenoScreen, Lille, France. From March 2015 to December 2015, the samples were sequenced on an Illumina NovaSeq 6000 (2 x 150 bp) at CNAG. About 19 billion reads were produced for SOLA. In total, ∼ 64 billion reads were produced for BBMO-SOLA.

For the surface global ocean, we used a metagenomic dataset generated during the *Tara Oceans* expedition 2009–2013^46,48^. This dataset comprises surface water samples collected from 82 stations from 2009 to 2013, encompassing the 0.2–1.6 and 0.22–3 μm size fractions (129 metagenomes; **Supplementary Dataset S2 and S3**). The two fractions were collected sequentially rather than concurrently: early-leg stations were filtered through 0.2–1.6 μm membranes, whereas later stations were processed with 0.22–3 μm filters. Hereafter, the three metagenomic datasets will be referred to as BBMO, SOLA, and TARA, respectively (**Figure S1**).

### Initial metagenome processing

All metagenomes were cleaned with *cutadapt 1.16*^54^ using a quality threshold of 20 for both 5’ and 3’ ends, a minimum length corresponding to half the size of the metagenomic read length, and Illumina-trueseq adapters (R1=AGATCGGAAGAGCACACGTCTGAACTCCAGTCA, R2=AGATCGGAAGAGCGTCGTGTAGGGAAAGAGTGT). TARA and SOLA metagenomes from January 2012 until February 2015 were sequenced with a read length of 100 base pairs (bp), while all BBMO and the rest of the SOLA metagenomes were sequenced with a read length of 150 bp. Lastly, all TARA metagenomes belonging to the same station were concatenated together for simpler downstream analyses.

### Metagenomic information content

Due to the high computational demands, the BBMO Metagenome-Assembled Genomes (MAGs) were delineated using a subset of the 15-year metagenomic dataset, comprising 7 years of monthly samples (January 2009 to December 2015), totaling 84 samples. We aimed at co-assembling groups of metagenomes with comparable information content. To define these groups, we determined pairwise metagenome similarity using Simka v1.5.2^55^, with a k-mer size of 21, a minimum Shannon Index of 1.5, a minimum read size of 70, and Bray-Curtis dissimilarities. We clustered the Bray Curtis distances using *hclust* and UPGMA in R^56^, which allowed us to define four groups (G) of samples corresponding to different times of the year: G1 (winter), G2 (spring), G3 (early summer), and G4 (late summer) including 37, 9, 14, and 22 metagenomes respectively (**Figure S2**). Two samples were excluded as they did not cluster into any group.

### Co-Assembly and reconstruction of Metagenome-Assembled Genomes (MAGs)

Samples from each cluster were co-assembled together using MegaHIT v1.2.8^57^ with preset *meta-large* and 750 gigabytes of RAM. Before the binning step, to obtain contig abundances across samples, the 84 BBMO metagenomes (years 2009–2015) were back-mapped to the four co-assemblies using BWA v.0.7.17-r118^58^ in default mode. Unmapped reads, secondary hits, and reads with an alignment quality below 10 were removed using Samtools^59^ v1.8 (for G2, G3, and G4) and v1.12 (for G1). The resulting BAM files were used as input to MetaBAT v2.12.1^60^, run in default mode with a minimum contig length of 2.5 kb, to generate four different sets of MAGs for each co-assembly. The contig depth values per metagenome from MetaBAT, along with the original BAM files, were provided as input to two other metagenomic binners to generate additional MAGs: Concoct v0.4.2^61^ and MaxBin2 v.2.2.5^62^; both ran in default mode using a minimum contig length of 2.5 kb. Then, to further improve the quality and accuracy of the predicted genomes, all MAGs from the three binners were combined and refined with the MetaWrap v1.3-4bf5f8a pipeline^63^ in default mode. Only refined MAGs with completeness ≥ 50% and contamination ≤ 10% were kept^64^. A total of 2,311 MAGs were obtained (909, 347, 457, and 598 MAGs for G1, G2, G3, and G4, respectively), which were taxonomically annotated with GTDB-Tk v1.5^65^ in default mode with the *classify_wf* workflow (reference database version r202). To remove redundancy, while keeping closely related MAGs, a dereplication was performed at 99% Average Nucleotide Identity (ANI) with dRep v2.3.2^66^ for all MAGs, independently of the original group (G). Ultimately, a total of 1,505 high-quality, non-redundant MAGs were generated (**Figure S3**, **Supplementary Dataset S4)**. Gene predictions and functional annotation were carried out using both Prokka v1.14.6^67^ and EnrichM v0.5.0’s database v10^68^.

### Abundance and Horizontal Coverage across samples

To obtain MAGs’ abundances across all datasets and to reduce the number of incorrectly mapped reads, we performed a competitive mapping approach. First, all 1,505 MAGs were concatenated into a single FASTA file. Second, all clean reads from TARA, BBMO, and SOLA were mapped with BWA-MEM2 v2.2.1^69^ against the concatenated file. Only reads with an identity greater than 95% and an alignment coverage greater than 80% were retained. Lastly, RPKM values (number of mapped reads per kilobase of genome per million mapped reads) and Genomic Horizontal Coverage (percentage of the genome covered by at least one filtered read) were calculated for each metagenomic sample using CoverM v0.7.0^70^.

### Variant calling and genetic differentiation

To assess the population-level variation of the prokaryotic MAGs through our spatiotemporal datasets, we predicted Single Nucleotide Variants (SNVs), insertions, and deletions across the three datasets. For each studied scale, temporal (BBMO + SOLA) and spatial (TARA) sets, all BAM files were merged into a single file with the Samtools 1.8 *merge* function. The merged BAMs were then given as input to Freebayes v1.3.1^71^ to perform Variant Calling for all MAGs combined with ploidy set to one (-p 1) and a minimum of four observations to support alternate alleles (-C 4). The generated variant call files (VCF) for each MAG were filtered to retain only the contigs of interest using BCFtools v1.19^72^. They were given to POGENOM v.0.8.4^73^, with a minimum coverage for a locus set to ten (--min_count 10) and a minimum number of stations for a locus to be present set to four (--min_found 4).

This analysis enabled the computation of several parameters for each MAG: 1) Nucleotide diversity (ρε), defined as the average number of nucleotide differences per site between any two randomly chosen sequence reads from the sample population; 2) The fixation index (F_ST_), which measures the extent to which the genetic diversity is structured among populations by estimating the proportion of total genetic diversity attributable to allele frequency differences across all pairwise comparisons between samples; and 3) The non-synonymous vs. synonymous mutation (pN/pS) ratios for each predicted gene across samples. To focus downstream analyses on candidates potentially under positive selection, we retained genes with a mean pN/pS ratio > 0.8 within each dataset (temporal or spatial).

Genomic populations were delineated by clustering genomes based on their pairwise genetic differentiation (F_ST_). Specifically, we computed an all-against-all F_ST_ matrix and performed hierarchical clustering in R (*hclust*^74^; UPGMA/average linkage), which progressively merges clusters using the average F_ST_ among all genome pairs across the two candidate clusters^75^. We refer to this cluster-to-cluster average F_ST_ as *Mean cF_ST_*. In addition, for each MAG we computed an overall mean differentiation value (*Mean F_ST_*) defined as the mean of all its pairwise F_ST_ values within a dataset.

A Permutational Multivariate Analysis of Variance (PERMANOVA) was carried out using the adonis2 function from the *vegan* (v2.6-10) R package^76^ to assess the percentage of the F_ST_ variance that changes in temperature, day length, salinity, PO_4_, NO_3_, NO_2_, and Si could explain. Environmental variables were previously z-score normalized with the *scale* function, and the model was run with by = “terms” (Type I sums of squares) and 9,999 permutations. Any samples with ‘NA’ in the F_ST_ matrix were excluded before analysis.

A Mantel test was conducted to assess the correlation between F_ST_ distance and both temporal (days) and geographical (kilometers) distances between samples. Marine least-cost path geographical distances between TARA stations were calculated using the *lc.dist* function from the *marmap* (v1.0.10) R package, ensuring connections were constrained to surface waters by setting a depth limit of 0 meters. The Mantel test was performed using the *mantel* function from the *vegan* (v2.6-10) R package. All analyses and visualizations were conducted using R (v4.4.0) (available through a Zenodo record; see Data availability section).

## RESULTS

### Stronger spatial than temporal population differentiation

We reconstructed 1,505 Metagenome-Assembled Genomes (MAGs) from the Blanes Bay Microbial Observatory (BBMO) long-term time series. On average, these genomes were of high quality, with a mean completeness of ∼79% and a mean contamination of ∼2.6%. The collection spans both Bacteria (1,444 MAGs) and Archaea (61 MAGs). At the phylum/class level, the most represented lineages were Gammaproteobacteria (373 MAGs), Alphaproteobacteria (339 MAGs), Bacteroidota (355 MAGs), Verrucomicrobiota (87 MAGs), Myxococcota (68 MAGs), and Thermoplasmatota (58 MAGs), together with additional contributions from Planctomycetota, Actinobacteriota, Cyanobacteria, and other lineages that are recurrent in marine planktonic communities (**Figure S3, Supplementary Dataset S4**). This broad taxonomic representation captured a moderate fraction of the microbial metagenomic diversity of the NW Mediterranean coast, accounting for 25.3% of the metagenomic reads at BBMO, 24.2% at SOLA, and 5.8% at the global scale.

We assessed population divergence (PD) in each of the 1,505 MAGs retrieved from BBMO across the two temporal datasets, BBMO and SOLA, and the global spatial dataset, TARA. Overall, PD was higher at the spatial scale than at the long temporal scale, while PD in BBMO and SOLA were very similar (**Figure 1A**). We classified PD into four categories based on the Fixation Index (F_ST_) values: Little PD (F_ST_ < 0.05), Moderate PD (0.05 ≤ F_ST_ < 0.15), High PD (0.15 ≤ F_ST_ < 0.25), and Very high PD (F_ST_ ≥ 0.25)^77^. In the temporal datasets, PD at BBMO was distributed as 25% Little, 45% Moderate, 15% High, and 15% Very High. At SOLA, the corresponding values were 25%, 47%, 15%, and 13% (**Figure 1B**). Overall, the proportions were comparable between the two temporal datasets, with BBMO showing a slightly higher proportion of Very High PD and SOLA a slightly higher proportion of Moderate PD.

**Figure 1.**
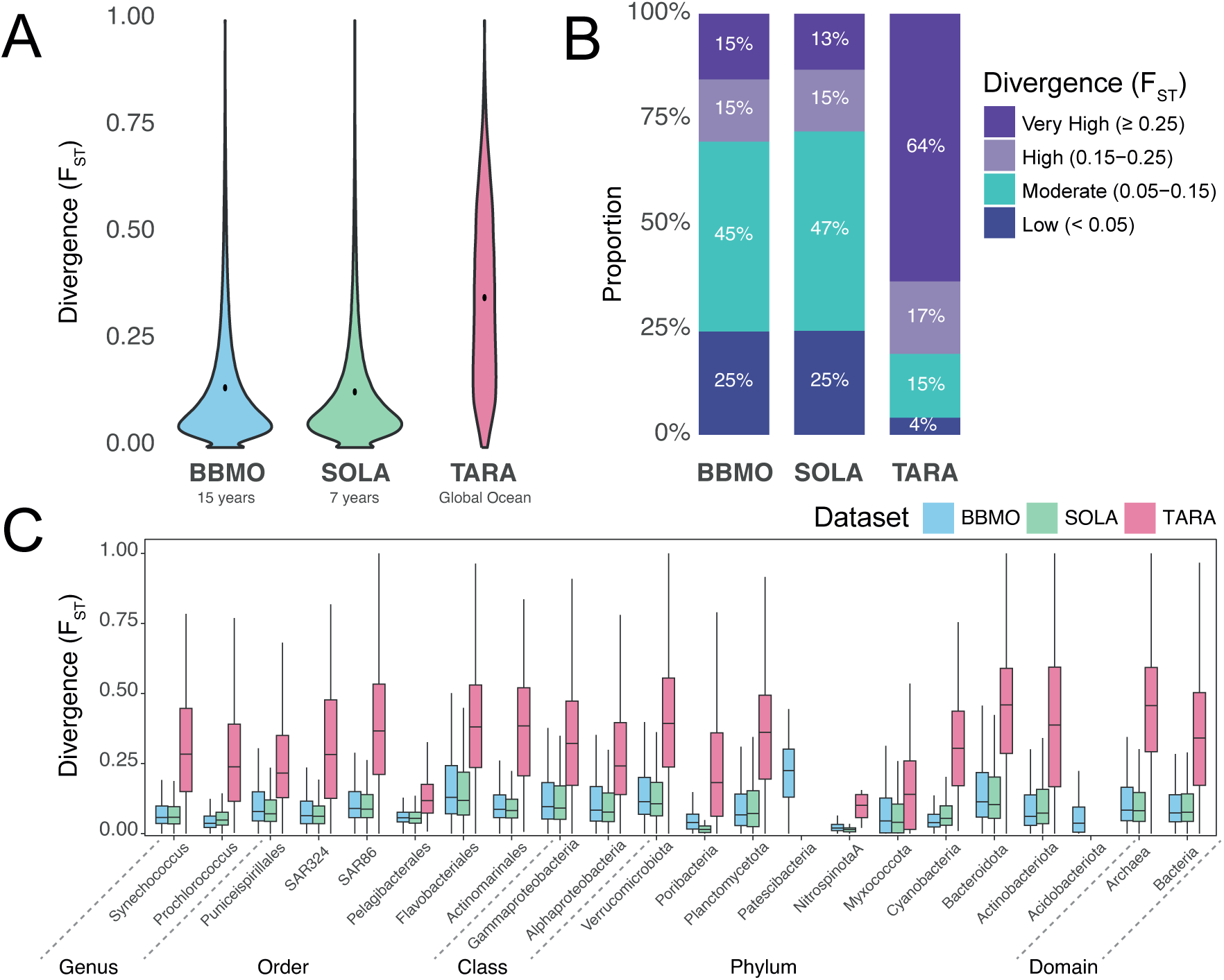
Population divergence of the 1,505 Mediterranean MAGs across temporal (BBMO and SOLA) and spatial (TARA; global ocean) scales. (A) Population divergence (F_ST_) over 15 and 7 years in the temporal datasets (BBMO and SOLA) and across the global ocean (TARA). (B) Distribution of F_ST_ values across four population divergence (PD) categories: Little PD (F_ST_ < 0.05), Moderate PD (0.05 ≤ F_ST_ < 0.15), High PD (0.15 ≤ F_ST_ < 0.25), and Very High PD (F_ST_ ≥ 0.25). (C) Distribution of population divergence (F_ST_) across taxonomic categories. Taxonomic ranks range from Genus to Domain, highlighting ecologically relevant groups that were selected as they represent abundant taxa in the temporal series or are groups with widespread global distributions. F_ST_ values were considered in the lowest taxonomic rank available only (*e.g.,* F_ST_ values within *Prochlorococcus* were not included in the Cyanobacteria or Bacteria categories). Only F_ST_ values from samples with at least 25% horizontal coverage of the corresponding MAG were considered. The temporal dataset spans up to 15 years of monthly surface samples, while the spatial dataset includes 82 surface stations from the global ocean (2009–2013) (Figure S1).

In contrast, in the global spatial dataset, PD was markedly higher, with 4% classified as Little PD, 15% as Moderate PD, 17% as High PD, and 64% as Very High PD (**Figure 1B**). These results indicate that, over a period of 7 to 15 years, the 1,505 MAGs from BBMO exhibited moderate population differentiation at the two neighboring Mediterranean locations but displayed a much higher degree of divergence across the global ocean. The same patterns were observed when PD was analyzed by taxonomic group (**Figure 1C**). However, on average, MAGs belonging to the *Pelagibacterales* (SAR11) order and the phyla Nitrospinota and Myxococcota exhibited lower PD in the global ocean than other taxonomic groups.

### Group-specific population differentiation

To ensure robust comparisons, we analyzed here only those genomes sufficiently represented across samples. Specifically, MAGs that had at least 25% genome coverage in a minimum of 35 out of the 265 temporal samples (BBMO and SOLA combined), and at least in 8 stations from the global ocean (TARA). In the 389 MAGs that met this criterion, we analyzed their population genomic differentiation patterns (**Supplementary Dataset S5)**.

Population divergence (PD) was classified as either weak (Mean F_ST_ < 0.15) or strong (Mean F_ST_ ≥ 0.15). Over 15 and 7 years, most MAGs exhibited weak PD (74.8%, 291 MAGs) in both time series, while only 25.2% (98 MAGs) displayed strong population differentiation. In turn, at the global ocean scale, strong PD was predominant (93.3%, 363 MAGs), with weak divergence being uncommon at this scale (6.7%, 26 MAGs). By integrating divergence patterns at both temporal and global scales, three main categories emerged: weak temporal–weak spatial (25 MAGs, 6.4%), weak temporal–strong spatial (266 MAGs, 68.4%), and strong temporal–strong spatial (97 MAGs, 24.9%). Only one MAG (G4.205, *Alcanivorax*) fell outside these categories, displaying strong temporal but weak spatial PD (**Figure 2A**).

**Figure 2.**
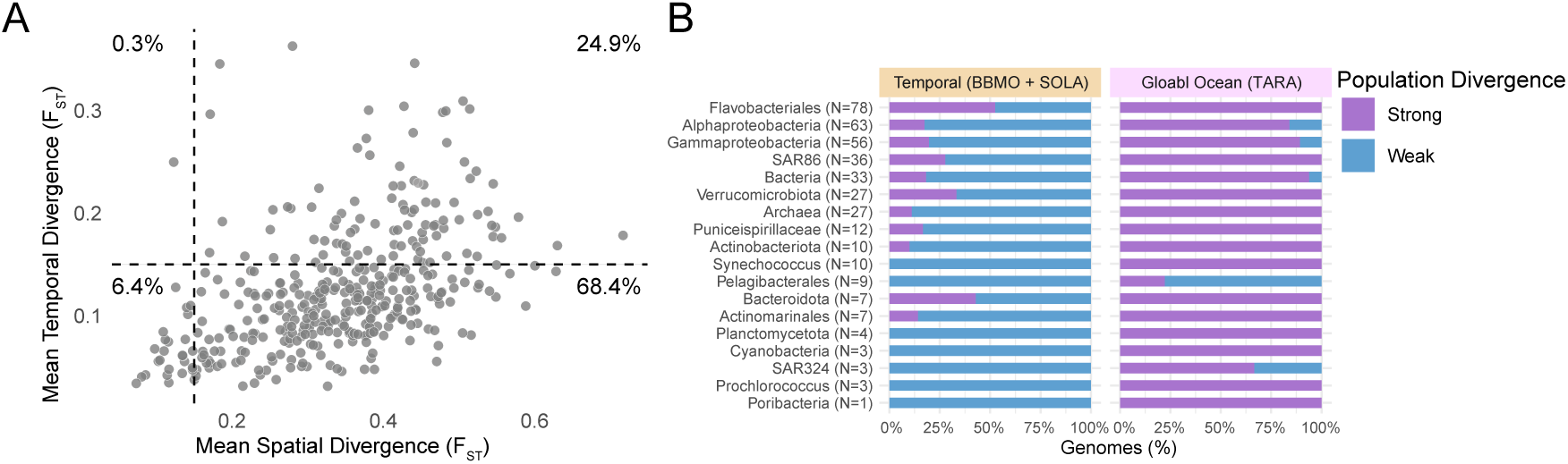
Population Divergence (PD) of Metagenome-Assembled Genomes (MAG). **(A)** Mean PD of each MAG in the temporal (BBMO [15 years] & SOLA [7 years]) and spatial (TARA [global ocean]) datasets. Each point represents one of the 389 MAGs that passed the filtering criteria (at least 25% of the genome covered in 35 and 8 samples from the temporal and spatial datasets, respectively). Dashed lines indicate the F_ST_ = 0.15 threshold, differentiating strong from weak PD. The percentages indicate the number of MAGs featuring strong or weak, spatial or temporal PD. **(B)** Proportion of genomes within each of the analyzed taxonomic ranks exhibiting strong or weak mean PD in each dataset. Taxonomic ranks range from Genus to Domain, highlighting ecologically relevant groups that were selected as they represent abundant taxa in the temporal series or are groups with widespread global distributions. Ranks are ordered based on the number of genomes (N) in each taxonomic group.

The predominant pattern of population divergence across the Mediterranean MAGs was characterized by weak temporal but strong spatial differentiation (68.4% of the MAGs). This points to the dominant role of large-scale spatial factors (*i.e.*, global ocean scale) in shaping population structure over time frames of 7–15 years at individual sites. This pattern was prevalent in abundant and ecologically relevant taxonomic groups, including *Prochlorococcus*, *Synechococcus*, *Actinomarinales,* and *Puniceispirillaceae* (SAR116 clade), all of which were consistently abundant throughout the temporal series. Additionally, the abundant SAR324 clade exhibited a mixed population divergence at the spatial scale, with one out of three MAGs exhibiting weak spatial and temporal differentiation (**Figure 2B**).

Specific taxonomic groups exhibited divergence patterns that contrasted with the overall trend. For instance, nearly all *Pelagibacterales* (SAR11) MAGs showed weak temporal divergence, with seven out of nine also exhibiting weak spatial divergence. In contrast, SAR86 and Flavobacteriales MAGs uniformly displayed strong spatial divergence but a notable heterogeneity in temporal divergence. Approximately 30% of SAR86 and nearly 50% of *Flavobacteriales* MAGs exhibited strong temporal divergence (**Figure 2B**), indicating significant variability in population differentiation within these groups across 7–15 years. These examples highlight the taxon-specific variability in population genomic differentiation and underscore the importance of examining divergence patterns for specific MAGs.

### Population divergence and structure

The Fixation Index (F_ST_) is widely used to assess population divergence, yet the thresholds for delineating populations remain ambiguous. Using cut-offs based on the literature^77^, we classified the overall mean population divergence (PD) as strong or weak based on a value of F_ST_ = 0.15. Furthermore, we explored population structure by clustering samples where taxa are present (*i.e.*, ≥ 25% of the genome covered) based on their pairwise F_ST_ distances.

Across the 389 MAGs, the number of clusters, used here as proxies for potential populations, decreased as the F_ST_ threshold increased. In the two time series, F_ST_ cutoffs of 0.10, 0.15, and 0.25 yielded 2,415, 1,149, and 557 clusters, respectively. By contrast, in the global ocean (TARA dataset), cluster counts fell more modestly, from 1,447 at F_ST_ = 0.10 to 1,429 at F_ST_ = 0.15 and 1,182 at F_ST_ = 0.25. This disparity implies that finer-scale population structure is more prevalent in the time series than at the global ocean scale. To further explore how population structure varies across lineages, we examined selected taxonomic groups of ecological relevance (graphs for all MAGs are available at Zenodo; see Data availability section).

### Limited temporal but pronounced spatial differentiation in Cyanobacteria

All MAGs from *Synechococcus* (10 MAGs) and *Prochlorococcus* (3 MAGs) exhibited the most common pattern among the studied genomes, characterized by weak temporal and strong spatial population divergence. In the temporal datasets, using a Mean cF_ST_ of 0.15 was insufficient to delineate genomic populations in most cases. For instance, in the *Prochlorococcus* MAG G4.203 (83.5% completeness, mean F_ST_ = 0.08), no potential populations were identified at this threshold. However, lowering the Mean cF_ST_ to 0.10 revealed five potential populations (**Figure 3A**). One population (T1) was exclusive to colder waters and had low abundance, while another (T2) consisted of only two samples from the warmer seasons. A third population (T3) was exclusive to the SOLA station, while a fourth (T4) comprised only a single BBMO sample. The largest and most abundant population (T5) exhibited the highest intradiversity, encompassing samples from both cold and warm waters across the two time series. Yet, seasonal structuring was evident within the group, as samples tended to cluster distinctly by thermal regime (**Figure 3A**). In total, 62.4% of the F_ST_ variance could be explained by the measured environmental factors in the time series, including temperature (39%), daylight hours (18.1%), and total chlorophyll a pigment (5.3%) (PERMANOVA, *p* < 0.05).

**Figure 3.**
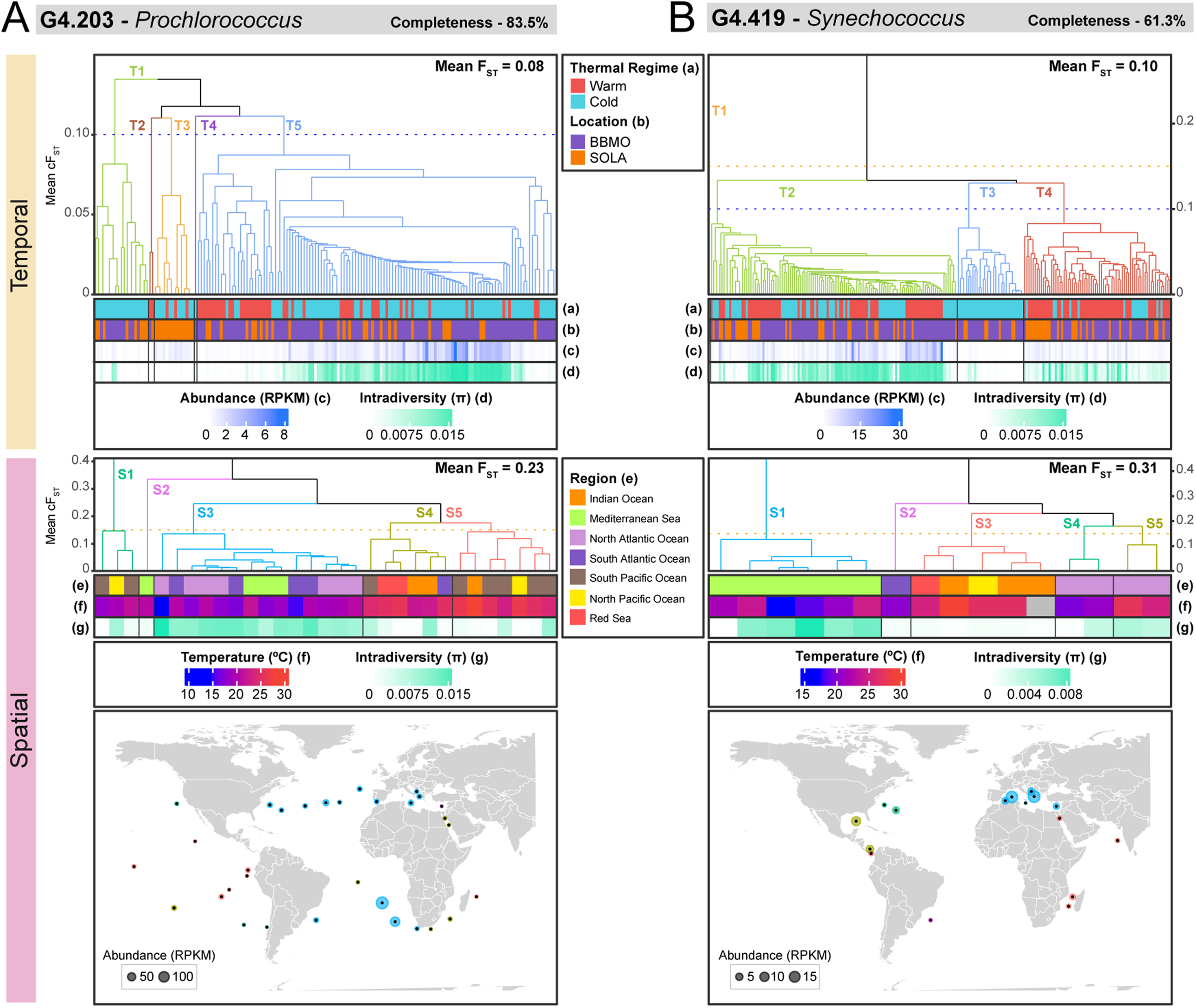
Population structure in the two time series (BBMO and SOLA) and the global ocean (TARA) for (A) *Prochlorococcus* MAG G4.203 and (B) *Synechococcus* MAG G4.419. Potential populations were defined using a Mean cF_ST_ threshold of 0.10, computed with the UPGMA clustering algorithm (dendrogram axis), which represents the average F_ST_ for each cluster. In turn, the Mean F_ST_ value indicates the overall mean of all pairwise F_ST_ values for each genome within each dataset. Different colors and labels distinguish between potential populations, labeled as T for Temporal and S for Spatial datasets. For each dataset, the main thermal regimes (Warm vs. Cold waters), location (BBMO vs. SOLA), abundance (RPKM, reads per kilobase of genome and million mapped reads), intradiversity (π), and temperature (°C) are displayed. Missing values are shown in grey. Genome completeness is also reported.

A similar pattern was observed in *Synechococcus* MAGs, where, for example, MAG G4.419 (61.3% completeness, mean F_ST_ = 0.10) exhibited four potential populations at Mean cF_ST_ = 0.10 (**Figure 3B**). The first potential population (T1) was represented by a highly divergent single sample (F_ST_ ∼ 0.2). The second potential population (T2) was the most abundant and diverse, encompassing samples from both time series and different seasons. The third population (T3) was only present in cold waters, while the fourth (T4) was mostly observed in warm water samples. The population structure appeared to be driven by environmental variation, with daylight hours (73.9%), temperature (4.2%), salinity (0.7%), and PO_4_ (1.1%) explaining 80% of the F_ST_ variance (PERMANOVA, *p* < 0.05).

In the global ocean, these *Prochlorococcus* and *Synechococcus* genomes exhibited high mean F_ST_ values (0.23 and 0.31, respectively), indicating the presence of highly divergent populations (**Figure 3**). Each genus displayed five potential populations in the analyzed global dataset, although they were structured differently. *Prochlorococcus* populations appeared to be primarily influenced by temperature, with potentially different populations in samples from colder subtropical waters (S1–3) versus those from warmer tropical regions (S4–5) (**Figure 3A**). *Prochlorococcus* exhibited a widespread geographic distribution and was most abundant in the South Atlantic Ocean, despite its representative MAG being initially recovered from the Mediterranean Sea (BBMO). In contrast, *Synechococcus* populations appeared to be structured by oceanic basins, with distinct potential populations identified in the Mediterranean Sea (S1), South Atlantic Ocean (S2), Indian Ocean (S3, including Red Sea and one North Pacific Ocean samples), and North Atlantic Ocean (S4–5) (**Figure 3B**). *Synechococcus* exhibited a higher abundance and intradiversity in the Mediterranean Sea (S1), and a less widespread geographic distribution than *Prochlorococcus*. Silicate (Si) explained 12.3% of the F_ST_ at the global scale in *Prochlorococcus* (PERMANOVA; *p-value < 0.05*). This association could be interpreted as correlational, as silicate covaries with other oceanographic and nutrient gradients at the global scale. None of the measured environmental variables accounted for any of the observed F_ST_ variation in *Synechococcus*. Thus, either population structuring was not directly explained by most of the measured environmental variables, or the limited sample size (31 and 16 samples in the global ocean for *Prochlorococcus* and *Synechococcus*, respectively) was insufficient to detect broad structuring patterns.

### Exceptions to the dominant pattern: SAR11 and SAR86

While *Prochlorococcus* and *Synechococcus* followed the most frequently observed pattern of weak temporal and strong spatial population divergence, not all taxa exhibited this trend (**Figure 2A**). Other MAGs showed distinct population divergence patterns, with either consistently weak or consistently strong divergence across temporal and spatial datasets. Members of the SAR11 and SAR86 clades, two highly abundant and ecologically relevant marine bacterial groups, exemplified these patterns. SAR11 MAGs displayed remarkably low levels of population divergence across both temporal and spatial scales. All nine genomes exhibited weak population divergence over 15 and 7 years in both time series, and most (7 out of 9) also displayed weak differentiation at the global ocean scale (**Figure 2B**). In contrast, SAR86 MAGs showed consistently high spatial population divergence, with all 36 genomes classified with strong PD at the global ocean scale. Temporal population divergence within SAR86 was variable, with approximately 30% of the genomes exhibiting strong PD over 7 and 15 years, while the rest displayed weak PD (**Figure 2B**).

Illustrative examples of the previous patterns include the SAR11 MAG G2.171 (66.7% completeness), which showed consistently weak population divergence across temporal and spatial datasets, and the SAR86 MAG G1.297 (70.8% completeness), which exhibited strong divergence at both temporal and spatial scales (**Figure 4**). The SAR11 MAG G2.171 had mean F_ST_ values of 0.08 (temporal) and 0.11 (spatial) (**Figure 4A**). Despite the low overall divergence, population structure was still detectable at a Mean cF_ST_ threshold of 0.10 in the temporal dataset and 0.15 in the spatial dataset (**Figure 4A**). Within the time series, four SAR11 MAG G2.171 populations were identified, all displaying high abundance and intradiversity, except for one (**Figure 4A**). T1 was exclusively composed of SOLA samples and exhibited the lowest intradiversity among all clusters. The other populations displayed thermal structuring, with T2 and T3 primarily composed of cold-water samples from both BBMO and SOLA (**Figure 4A**). On the contrary, T4 was associated with warm-water samples. In total, 47.4% of the F_ST_ variance in the temporal dataset could be explained by the measured environmental factors, such as temperature (44.3%), daylight hours (1.5%), and salinity (1.6%) (PERMANOVA, *p-value* < 0.05).

**Figure 4.**
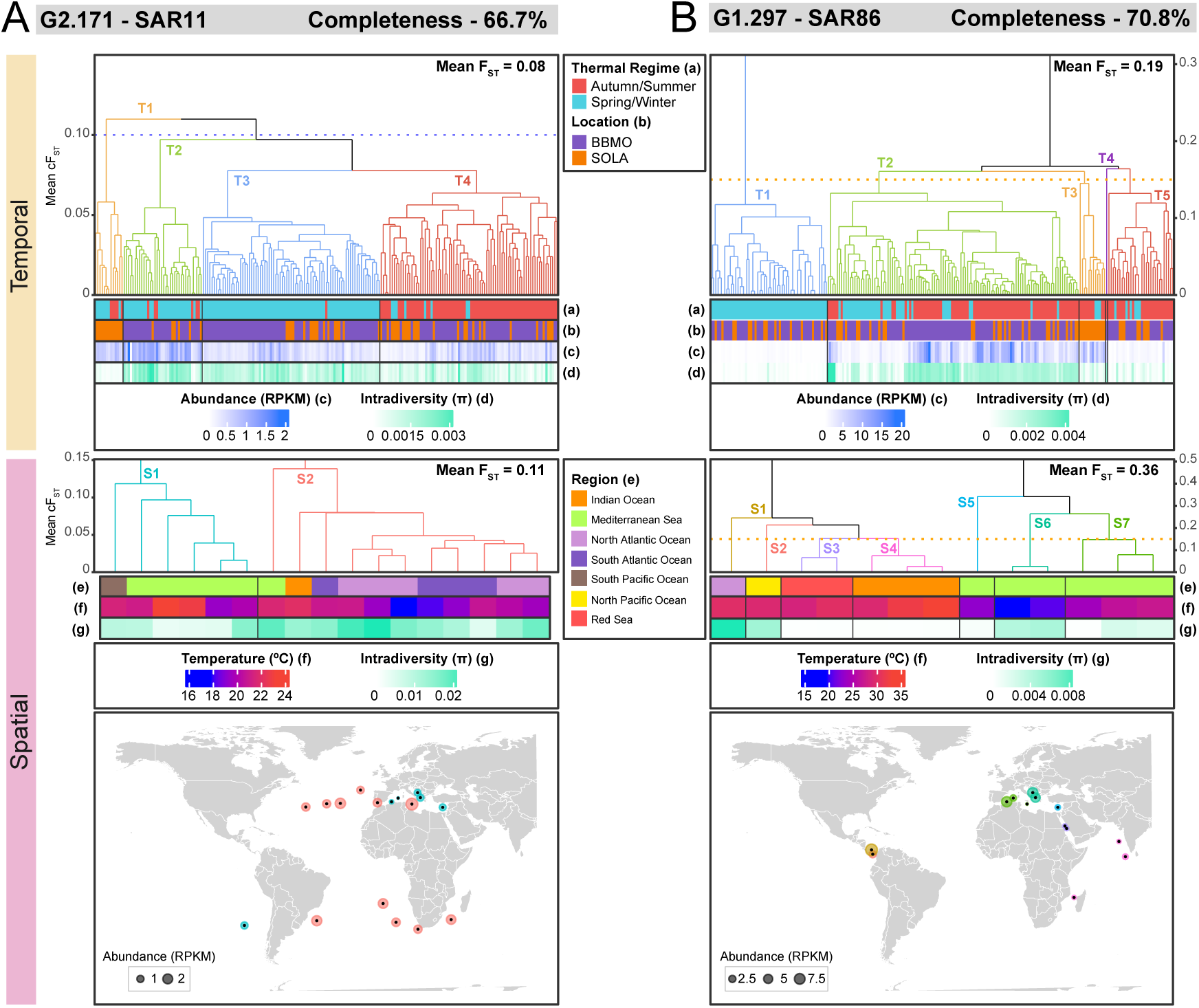
Population structure in the two time series (BBMO and SOLA) and the global ocean (TARA) for (A) SAR11 MAG G2.171 and (B) SAR86 MAG G1.297. Potential populations (clusters) were defined using thresholds of Mean cF_ST_ = 0.10 and 0.15 for SAR11 and SAR86 genomes, respectively (dendrogram axis), which represent the average F_ST_ for each cluster. The Mean F_ST_ value indicates the overall mean of all pairwise F_ST_ values for each genome within each dataset. Different colors and labels distinguish between potential populations, labeled as T for Temporal and S for Spatial datasets. For each dataset, thermal regime (Warm vs. Cold waters), location (BBMO vs. SOLA), abundance (RPKM, reads per kilobase of genome and million mapped reads), intradiversity (π), and temperature (°C) are displayed. Genome completeness is also reported.

In the global ocean dataset, two potential populations of the SAR11 MAG G2.171 were identified. S1 consisted mostly of Mediterranean Sea samples and exhibited lower intradiversity, whereas S2 included other subtropical samples and a higher intradiversity (**Figure 4A)**. The observed spatial structuring was not explained by the measured environmental variables (PERMANOVA; *p-value* > 0.05), suggesting that other variables may be driving population structure or that the limited sample size was insufficient to detect prevalent patterns. Overall, SAR11 MAG G2.171 displayed a stable, albeit low, abundance across all potential populations and samples.

In contrast to SAR11, SAR86 MAG G1.297 displayed strong population divergence at both temporal and spatial scales. It exhibited a mean F_ST_ of 0.19 over 15 and 7 years and 0.36 in the global ocean (**Figure 4B**). Five potential populations were identified in the time series using a Mean cF_ST_ = 0.15, each with distinct abundance and thermal preferences. T1 was present in cold-water samples from both SOLA and BBMO, displaying both low abundance and low intradiversity. In turn, T2 was the most abundant potential population and was detected in cold- and warm-water samples from both time series (**Figure 4B**). The internal structure of T2 suggested further differentiation at lower F_ST_ thresholds, indicating the presence of distinct subpopulations. T3 was exclusive to SOLA and primarily present in warm-water samples, while T4 was detected in a single BBMO sample. T5 was present in both time series, being mostly associated with warm waters, and exhibited low abundance and moderate intradiversity. Most of the F_ST_ variance could be explained by temperature (59.5%), while daylight hours (7.6%), total chlorophyll *a* (2.4%), and NO_2_ (2.7%) (PERMANOVA; *p-value* < 0.05), explained lower proportions.

In the global ocean, seven potential populations were identified for SAR86 MAG G1.297 (**Figure 4B**). S1 to S4 were detected in tropical regions, with S1 and S2 being the most abundant and exhibiting the highest intradiversity, particularly in the Atlantic population (S1). In contrast, S5 to S7 appeared exclusive to the Mediterranean Sea, with a structuring potentially linked to latitude and longitude. SAR86 G1.297 populations were most abundant in the Mediterranean Sea, consistent with the origin of this MAG from the BBMO. Among these potential populations, S6, present in the northeastern Mediterranean, exhibited the highest intradiversity. S5 occurred in the southeastern Mediterranean, close to the Red Sea, whereas S7 was found in the western Mediterranean, the region from which the SAR86 MAG G1.297 originated. None of the environmental variables measured could explain the spatial variance in F_ST_ (PERMANOVA; *p-value* > 0.05). In sum, despite its limited abundance in most oceanic regions, SAR86 G1.297 exhibited differentiated population structure in the Mediterranean Sea, where it was comparatively more abundant.

### Patterns of positive selection across temporal and spatial scales

We evaluated the potential influence of positive selection across the 389 MAGs to determine if adaptive processes could explain the observed population differentiation. To do so, we calculated the ratio of non-synonymous to synonymous mutations (pN/pS) for all genes within each MAG in both the temporal (BBMO and SOLA) and spatial (global ocean) datasets. We focused specifically on genes with a mean pN/pS greater than 0.8. Interestingly, the number of positively selected genes per MAG was higher in the temporal datasets compared to the global ocean (Wilcoxon signed-rank test [paired, one-sided]; *p-value* < 0.01), despite the latter showing higher overall population divergence (**Figure S4A**). This trend was consistent when analyzing positively selected genes across taxonomic groups (**Figure S4B**). The taxa with the highest average number of positively selected genes per MAG in both temporal and spatial datasets included Gamma- and Alphaproteobacteria, as well as the SAR324 and SAR116 (Puniceispirillales) clades (**Supplementary ataset S6**).

We also identified a positive correlation between genome size and the total number of positively selected genes in both the temporal and spatial datasets (**Figure S4C**). After normalizing by genome size (genes per megabasepairs; Mbp), specific taxonomic groups showed elevated positive selection. In the time series, MAGs from the orders Flavobacteriales and SAR86 (*e.g.,* MAG G1.297, **Figure 4B**), as well as the classes Alpha- and Gammaproteobacteria, displayed the highest density of positively selected genes (**Supplementary Dataset S6**). In the global ocean dataset, the highest density of positively selected genes per Mbp occurred in the class Gammaproteobacteria, genus *Synechococcus* (*e.g.,* MAG G4.419, **Figure 5**), and orders Flavobacteriales and Puniceispirillales.

**Figure 5.**
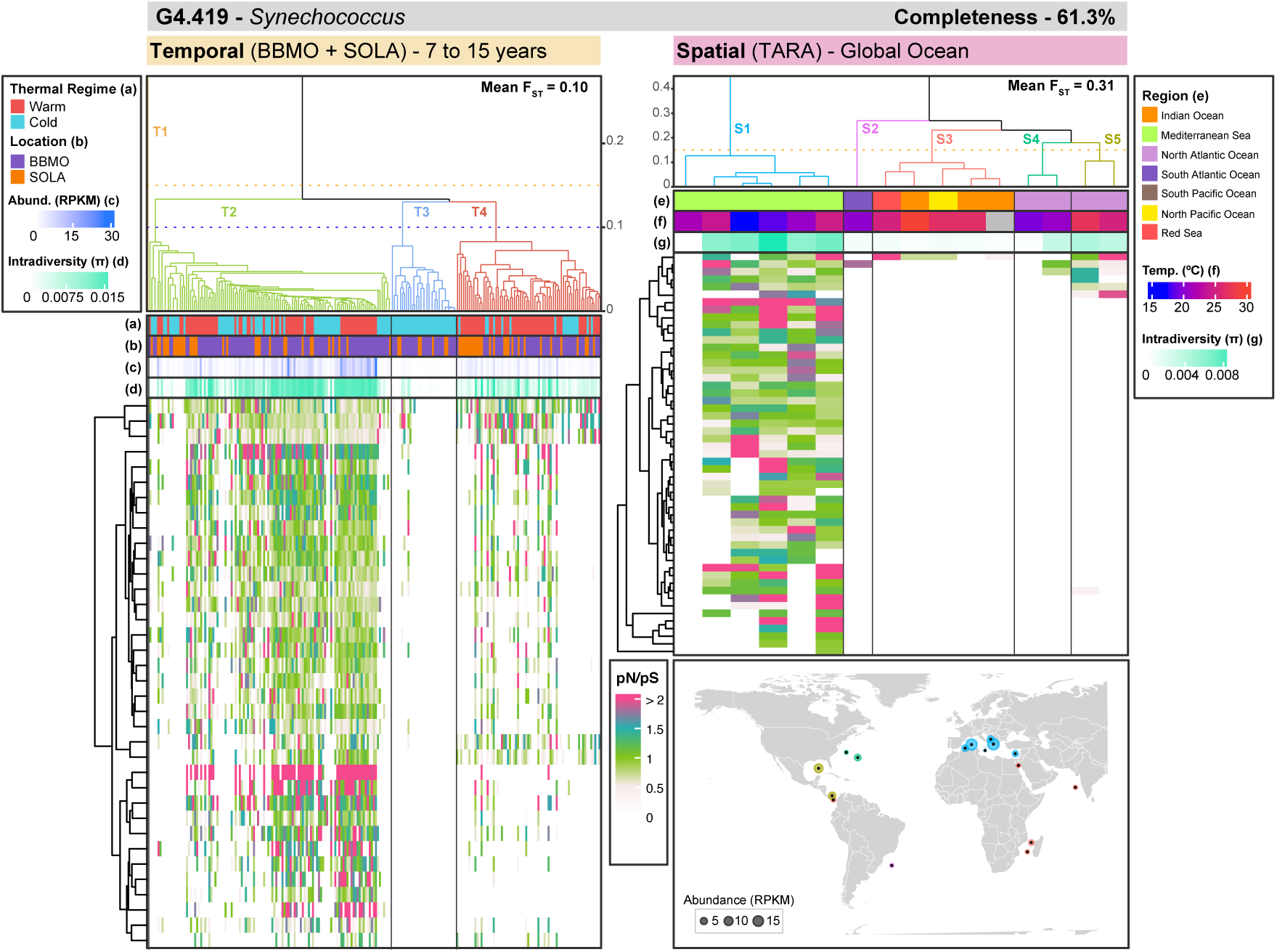
pN/pS values for positively selected genes (mean pN/pS > 0.8) across samples and populations of *Synechococcus* MAG G4.419, analyzed at both temporal (BBMO and SOLA) and spatial (global ocean) scales. The cell colors in the heatmaps indicate the pN/pS of a positively selected gene in a given sample. White tails indicate NA values (pS = 0 in pN/pS calculations), which may occur either due to limited mapping or the absence of synonymous variants. The mean pN/pS is computed omitting NA values. Clustering is based on the F_ST_ distances between samples. Clusters are defined using a mean cF_ST_ of 0.10 and 0.15 for the temporal and spatial datasets, respectively. The clusters, colors, and letters match those of the populations in Figure 3B.

A closer examination of the distribution of positively selected genes within specific genomes provided further insights into adaptive patterns (graphs for all MAGs are available at Zenodo; see Data availability section). For instance, the pN/pS analysis of *Synechococcus* MAG G4.419 (**Figure 5**) revealed distinct gene-level selection signatures closely aligned with the F_ST_-based population structure. Specific genes consistently showed elevated pN/pS ratios, indicating positive selection within particular samples. Higher pN/pS values were observed in samples characterized by higher genome-wide intradiversity and greater abundance (**Figure 5**). This suggests that the ability to detect positive selection may depend on population abundance in the investigated samples, with near-zero pN/pS values in genes from low-abundance samples potentially reflecting limited detection rather than an actual absence of selection.

In the time series (BBMO and SOLA), the pN/pS ratios of positively selected genes in *Synechococcus* varied among potential populations, reflecting possible population-specific adaptations to different seasons or niches (**Figure 5**). For instance, the *Synechococcus* potential population T2, comprising both cold and warm water samples, exhibited the highest pN/pS values in the analyzed positively selected genes, pointing to multiple population-specific adaptations. Similarly, population T4, which was predominantly associated with warm waters, also exhibited multiple positively selected genes, indicating potential population-level adaptations (**Figure 5**). Both T2 and T4 also displayed elevated genome-wide intradiversity. In contrast, population T3, exclusively present in cold-water samples and characterized by low abundance, showed minimal evidence of positive selection in the analyzed genes (**Figure 5**). These results suggest the presence of ecologically distinct *Synechococcus* populations with specific adaptive traits across the 15- and 7-year time series.

In the spatial dataset, the pN/pS ratios of positively selected genes exhibited a clear contrast between Mediterranean samples and those from other basins (**Figure 5**). Mediterranean samples exhibited substantial variability in the pN/pS ratios of positively selected genes, accompanied by clear signals of positive selection, whereas most samples from other oceanic basins showed no comparable evidence (**Figure 5**). Since all Mediterranean samples belonged to population S1, the signals of positive selection likely reflect adaptations specific to this population. The potential populations from the tropical (S3) and subtropical regions (S2, and S4–5) displayed limited evidence of positive selection in the analyzed genes (**Figure 5**). The genome-wide intradiversity also displayed variation, with populations S1 (Mediterranean), S5, and S4 (North Atlantic) displaying higher values than the other populations.

### Population differentiation as a function of temporal and geographic distances

We examined whether population differentiation (F_ST_) correlated with temporal or geographic distances. Correlations between F_ST_ and distance matrices (using either temporal [in days] or geographic [in kilometers] distances) were investigated across the 389 MAGs using Mantel tests. Significant (*p-value* < 0.05) positive correlations were observed for 154 (40%), 137 (35%), and 255 (66%) MAGs at both temporal (BBMO and SOLA) and spatial (TARA) scales, respectively (**Figure 6A, Supplementary Dataset S7**). This suggests that, in several MAGs, population divergence tends to increase as samples become more temporally or spatially distant from one another. However, Mantel correlation coefficients (r-values) were consistently higher for spatial than temporal datasets, suggesting that large geographic distances (*i.e.*, thousands of km) exert a stronger influence on population differentiation than long periods of time (*i.e.*, up to 15 years) (**Figure 6A**).

**Figure 6.**
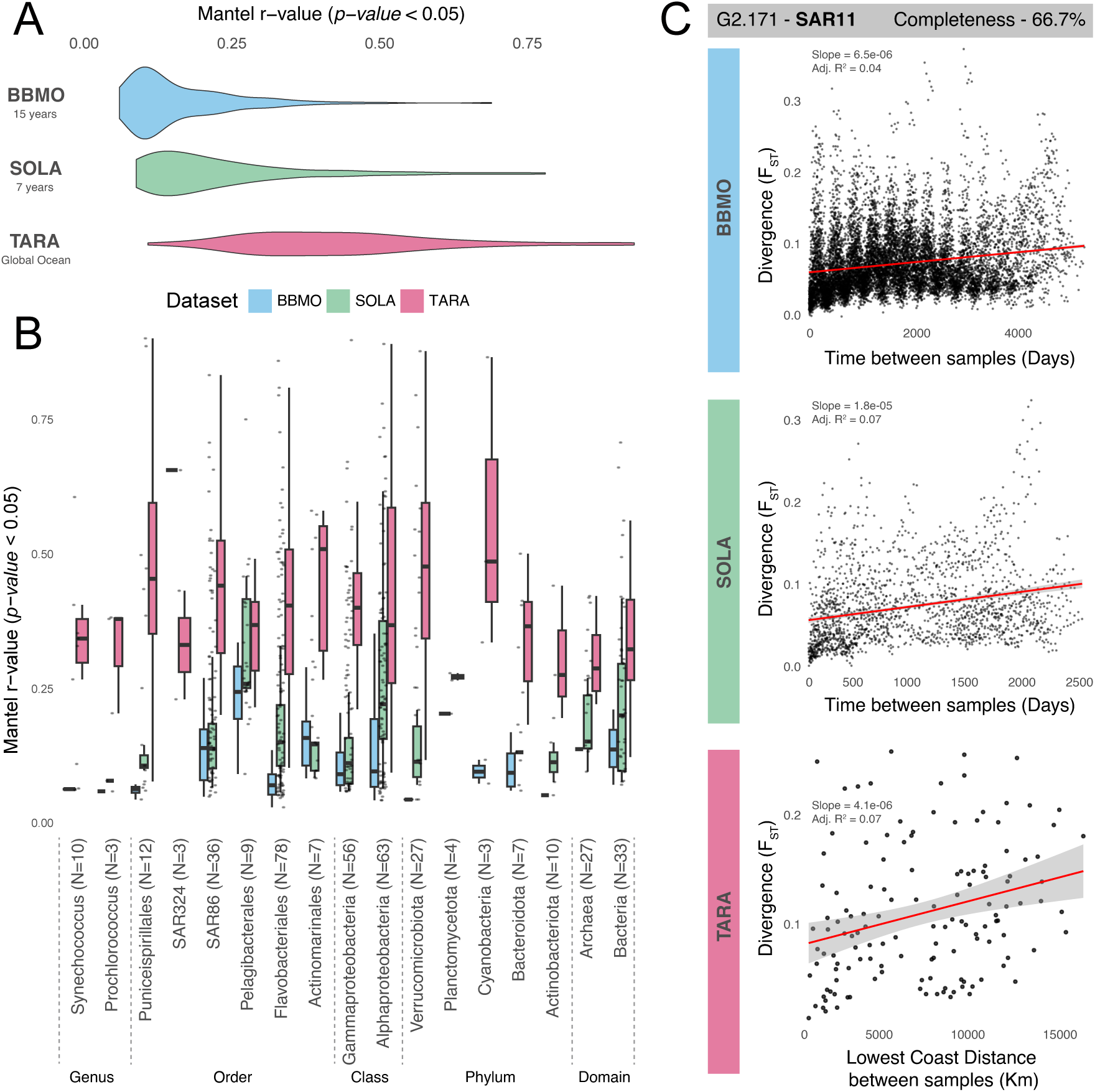
Population divergence (F_ST_) vs. temporal or geographic distance. (**A**) Distribution of significant Mantel correlation coefficients (r-values, *p-value* < 0.05) between pairwise population divergence (F_ST_) matrices and temporal (days; BBMO and SOLA datasets) or geographic (km; TARA dataset) distances for the 389 MAGs analyzed. (**B**) Mantel correlation coefficients grouped by taxonomic category for each dataset. Taxonomic categories are shown at various levels (genus to domain). The variable N indicates the number of MAGs considered in each taxonomical group. (**C**) Example illustrating population divergence patterns for MAG G2.171 (*Pelagibacterales*, SAR11 clade). The temporal datasets exhibit divergence patterns over time (days) in BBMO and SOLA, whereas the spatial dataset (TARA) displays divergence relative to geographic distances (km) between sampling locations. Points represent individual pairwise comparisons between samples, and trend lines illustrate the correlation between F_ST_ and temporal or geographic distance. The completeness of MAG G2.171 was 66.7%.

The temporal datasets revealed a dominant pattern of pronounced seasonal structuring in most MAGs (369 out of 389, 95%), characterized by reduced population differentiation among samples collected in the same season across years and increased differentiation between samples from opposite seasons (see example in **Figure 6C**). In addition, all MAGs with a significant Mantel correlation (*p-value* < 0.05; total 154 in BBMO, 139 in SOLA, and 81 in both simultaneously) exhibited a subtle yet consistent temporal trend of increasing population divergence with time, suggesting a gradual accumulation of genetic differentiation over the studied periods (7 to 15 years) (**Supplementary Dataset S7)**. This temporal divergence was particularly pronounced in the *Pelagibacterales* (SAR11 clade), which showed one of the strongest temporal correlations among all taxonomic groups analyzed (**Figure 6B**). Specifically, SAR11 MAG G2.171 exemplified this scenario: in the BBMO, a distinct seasonal pattern was observed alongside a slight increase in divergence over time, whereas in SOLA, seasonality was less evident, though the positive temporal correlation persisted (**Figure 6C**).

At the global ocean scale (TARA), correlations between population divergence and geographic distance were evident and consistent across most taxonomic groups (**Figure 6B**). Again, SAR11 MAG G2.171 clearly illustrated this spatial trend, showing a pronounced increase in population divergence with greater geographic separation between samples (**Figure 6C**). Thus, geographic distance appears to be a significant factor influencing the divergence of prokaryotic populations across large oceanic scales.

## DISCUSSION

Understanding how microbial populations vary genetically across space and time is key to elucidating the structure, function, and adaptive potential of the ocean microbiome. Yet, such information remains scarce for most microbial species. Our work is a step forward in understanding the genetic diversity and structure of marine prokaryotic populations and in identifying the genomic basis of population adaptation. We examined the population-level genomic differentiation of 1,505 Mediterranean genomes over extensive temporal (7 to 15 years) and geographic scales (global ocean). Although these MAGs represent only a fraction of total diversity, they encompass a broad taxonomic range and likely capture population-level dynamics representative of many other marine microbes across extended spatiotemporal scales.

The F_ST_ index was similar for the 1,505 MAGs in the two time series, encompassing 15 (BBMO) and 7 (SOLA) years, indicating that populations at these two locations in the Northwestern Mediterranean Sea, separated by ∼ 150 km, exhibit comparable levels of genetic differentiation. In fact, samples containing members of the same potential populations were detected in both time series, suggesting that the two locations share specific populations. Most genomes exhibited significant seasonal abundance patterns, with potential populations recurring annually. Microbial communities in both BBMO and SOLA, investigated using metabarcoding of the rRNA gene, revealed strong seasonal trends^43–45,78–82^, yet whether the same populations inhabited both sites had never been tested. Our results align with previous observations indicating that oceanic patch sizes, composed of relatively homogeneous microbial communities, range from a few to tens of kilometers^83,84^.

Nevertheless, differences in population differentiation between BBMO and SOLA were also observed, which may reflect seasonal variations in environmental conditions driven by the distinct climatological characteristics of each site. For instance, SOLA experiences occasional strong winter storms that bring nutrients from sediments into the water column and freshwater inputs from nearby rivers during flooding^43,78,85^. Additionally, differences in the number of samples between datasets covering distinct periods (BBMO: January 2008 to December 2022; SOLA: January 2009 to December 2015) might have contributed to variation in the F_ST_ estimates, although this is expected to be minor.

Population differentiation across MAGs at the global ocean scale was greater than the temporal differentiation observed within each long time series. To some extent, this may reflect the higher variability of surface-ocean environmental conditions across broad geographic scales compared to the seasonal variations observed at the two temperate time series in the Mediterranean Sea. Accordingly, broader ranges were recorded for several environmental variables in the *Tara Oceans* survey compared with those measured in the BBMO–SOLA time series (temperature: 32.0 vs. 18.0 °C; salinity: 17.0 vs. 4.0 PSU; PO₄: 2.0 vs. 0.3 µM; NO₃: 29.7 vs. 10.4 µM, respectively, **Supplementary Dataset S1 & S3)**. Furthermore, consistent with the idea that microbial communities inhabit oceanic patches spanning a few to tens of kilometers^83,84^, we found a higher population differentiation at hundreds of kilometers within the Mediterranean Sea (TARA stations 7, 9, 18, 23, and 25) than across the measured temporal scales at the two Mediterranean time series. Similar spatiotemporal patterns were reported in the Baltic Sea, where genomic differentiation among bacterioplankton populations was also more pronounced over spatial distances (spanning a 1,700 km transect) than across two years of seasonal sampling at the Linneaus Microbial Observatory. In that study, pronounced environmental gradients, most notably salinity, were likely the main drivers of spatial differentiation, shaping population structure more strongly than temporal factors^73^.

Altogether, among the 389 MAGs that were well represented in the time series and the global ocean, we identified three main patterns of population structure based on the strength of genomic differentiation (mean F_ST_ ≥ 0.15 indicating strong differentiation; mean F_ST_ < 0.15 indicating weak differentiation): (i) strong differentiation across both temporal and spatial scales, (ii) weak differentiation across both, and (iii) weak temporal but strong spatial differentiation. These patterns are consistent with the recognition that both geographic and environmental factors shape population structure in marine microbial communities^86^.

In general, for the analyzed MAGs originating from the Mediterranean Sea, pattern (iii) was the most common (68.4% of MAGs), while (i) and (ii) were specific to some genomes (24.9% and 6.4% of MAGs, respectively). Similar dominance of spatial over temporal differentiation has been observed in microbial communities of the Eastern Mediterranean using amplicon data, where community composition showed stronger associations with spatial gradients and water mass structure than with seasonal change^87^.

These findings suggest that large-scale spatial environmental gradients and geographic distance generally exert a stronger influence on microbial population composition than long-term seasonal variability. This has been reported by metagenomic studies showing that genomic variation in marine protists often aligns with geographic distance or environmental isolation, a process known as isolation-by-environment or isolation-by-distance^86^.

We detected only one instance of strong temporal but weak spatial differentiation among the analyzed genomes, suggesting that this pattern is rare across broad spatiotemporal scales. Populations of this MAG (G4.205, *Alcanivorax*) may be subject to strong seasonal selection, while experiencing limited spatial divergence, possibly due to high dispersal capacity or ecological generalism. To our knowledge, this pattern, *i.e.*, temporal genomic turnover outweighs spatial differentiation, has rarely been reported in marine bacteria. Most global-scale studies instead document either strong differentiation across both temporal and spatial axes^88^, or predominant spatial structuring with temporal stability^73^. The genomic dynamics of *Alcanivorax* may thus reflect genome-specific ecological strategies or niche shifts that are uncoupled from geographic distance, highlighting the complexity of microbial population responses to environmental fluctuations.

Clear examples of weak temporal, yet strong spatial differentiation within a single taxonomic group were observed among several *Prochlorococcus* and *Synechococcus* MAGs. Both taxa exhibited weak population differentiation over 15 and 7 years; however, low-divergence sub-populations (cF_ST_ = 0.10) could still be inferred from the F_ST_ data. Temperature and daylight accounted for a significant portion of the temporal variability in population differentiation in these MAGs. At the global ocean scale, although our analyses did not yield statistically significant results, we observed patterns indicative of potential adaptations to temperature and oceanic basins, in line with previous studies. Yan *et al.* demonstrated that *Prochlorococcus* high-light-adapted Clade II undergoes subclade differentiation linked to oceanographic features such as sea surface temperature, nutrient availability, and mixed layer depth, emphasizing the role of local adaptation in shaping population structure^37^. Similarly, Kent *et al.* found that both *Prochlorococcus* and *Synechococcus* follow parallel phylogeographic patterns across ocean basins, with temperature and nutrient availability driving genomic differentiation at large scales^38^. Furthermore, Doré *et al.* found that *Synechococcus* populations were structured along temperature gradients^89^, while Sharpe *et al.* identified genomic adaptations to local nutrient availability^90^.

Specific archaea and other bacterial groups also followed the general pattern of weak genomic differentiation over time but strong differentiation across the global ocean. In both time series, archaeal populations exhibited seasonal dynamics, as shown in the past both in SOLA, using MAGs for Ca. Poseidoniales, and in BBMO using 16S rRNA gene amplicon data^44,91^. Despite clear temporal variation in relative abundance, limited long-term population divergence was observed at both sites. A spatial structuring of populations was apparent across the global ocean for the MAG G3.122 — Nitrosopumilaceae, although it was not statistically significant (**Figure S5**). These findings align with previous evidence showing that specific archaeal distributions follow environmental gradients such as temperature, salinity, and nutrient availability, with distinct Thaumarchaeota clades and ammonia-oxidizing archaea occupying specific ecological niches shaped by physicochemical conditions^92^. A comparable pattern was found for SAR324 (Deltaproteobacteria), which in our dataset (2 MAGs) exhibited weak temporal differentiation but pronounced spatial structuring. This aligns with observations from the ALOHA time series, where genomic divergence remained low over time despite the presence of distinct ecotypes with depth- and season-specific distributions^93^.

Although most Mediterranean MAGs (74.8%) displayed weak population differentiation over 15 and 7 years, several others (24.9%) showed highly differentiated populations in both spatial and temporal scales. Such patterns were observed in ∼50% of the Flavobacteriales genomes. Studies based on the 16S rRNA gene indicate that the Flavobacteriales comprise diverse organisms inhabiting both warm, oligotrophic waters and cold, nutrient-rich waters in the North Atlantic, with apparent population-level differences between these environments^94–96^. Our population genomics analyses also showed distinct abundance patterns and strong genomic differentiation between cold and warm seasons at BBMO and SOLA, as well as across subtropical to subpolar regions globally. A good example was the Flavobacteriales genome G4.480 (**Figure S6**), which displayed a strong population structure driven by temperature (51.1%) and daylight hours (18.8%) in both time series, as well as by temperature (14.2%) and salinity (10.2%) in the global ocean (PERMANOVA, *p-value* < 0.05). This is consistent with observations from the Baltic Sea, where salinity has been shown to drive the differentiation of a Flavobacteriales population across its pronounced salinity gradient^73^. In our work, the 97 Flavobacteriales MAGs showing strong differentiation in both temporal and spatial scales suggest that some taxa experience comparable disruptive selection across broad spatiotemporal scales.

We found 25 MAGs (out of 389, 6.4%) that displayed weak population differentiation (Mean F_ST_ < 0.15) in both the time series and the global ocean. These cases represent exceptions to the general trend. Among them, SAR11 stands out. Despite their ubiquity and dominance in marine surface waters, SAR11 populations exhibited low genomic differentiation in both time series and across the global ocean. The large population sizes of SAR11^39^ may contribute to high dispersal rates, which, coupled with high levels of intra- and interspecies recombination^39^, could homogenize populations, limiting their divergence. Consistent with this, intrapopulation diversity in the analyzed SAR11 remained stable across both temporal and spatial scales, supporting well-mixed populations. Despite limited overall differentiation, SAR11 populations showed temperature-associated sub-structuring in both time series and possibly at the global scale. This aligns with earlier evidence that temperature drives population divergence in SAR11, reflecting fine-scale adaptations^35^. In contrast, Actinomarinales (7 MAGs), which overlap with SAR11 in terms of habitat and distribution^97,98^, displayed higher population differentiation, specifically at the global ocean scale. This pattern may be explained by their smaller population sizes^97^, which may limit dispersal and, in turn, promote population differentiation.

To assess whether the observed population differentiation results from adaptive processes, we evaluated signatures of positive selection in 389 genomes by calculating the ratio of non-synonymous to synonymous mutations (pN/pS) across all genes in each MAG, in both the temporal (15 and 7 years) and the spatial (global ocean) datasets. We found that the proportion of positively selected genes was higher in the long time series than in the global ocean, even though the latter exhibited greater overall population differentiation. This suggests that differentially adapted ecotypes are more apparent over longer temporal scales (up to 15 years in the time series) than over shorter ones (< 4 years in the global ocean), regardless of the spatial scale considered. Furthermore, this hints that a greater fraction of population differentiation in the global ocean may result from non-adaptive processes, such as genetic drift and isolation by geographic distance, compared with the time series^22^.

Typically, the number of positively selected genes per MAG was below 100 (**Figure S4**). This pattern is expected and indicates that positive selection targeted specific genes, fine-tuning them to slightly different ecological niches and potentially forming the basis of population-level adaptation. Comparable patterns have been reported in *Synechococcus* and *Sulfurovum*, where positive selection tends to be limited to a small subset of genes involved in nutrient acquisition and stress response in seasonally dynamic coastal waters (*Synechococcus*)^99^, or in phosphate limitation and chemical gradients in hydrothermal vent systems (*Sulfurovum*)^100^.

Groups such as Flavobacteriales, SAR116, SAR86, SAR324, Alpha- and Gammaproteobacteria consistently showed higher proportions of positively selected genes. This may reflect either a combination of biological signals, such as a higher rate of adaptive mutations in these groups, or technical factors like their overrepresentation in our MAG collection, which increases the detection power of selection. By contrast, SAR11 exhibited low signals of positive selection in the analyzed genes. Yet, recent analyses of the SAR11 pangenome suggest that population-level differentiation in this lineage may be driven by variation in the accessory genome rather than in ortholog mutations. Differences in gene content, particularly in ecologically relevant functions, have been shown to separate closely related SAR11 ecotypes with little core genome divergence^101^. This mechanism may extend to other highly abundant and cosmopolitan marine microorganisms that also exhibit weak population differentiation over time and space. In these species, low intrapopulation diversity and strong purifying selection could limit the accumulation of adaptive mutations, while differentiation may instead proceed via gene gain/loss dynamics in the non-core genome^102,103^.

Finally, we examined whether population differentiation scaled with temporal or geographic distance. A substantial fraction of MAGs showed significant positive correlations between F_ST_ and both temporal (40% and 35% of MAGs in BBMO and SOLA, respectively) and spatial distances (66% in TARA), indicating that populations become increasingly divergent over time and with geographic separation. However, spatial correlations were stronger than temporal ones, suggesting that large-scale geographic separation has a greater impact on microbial population divergence than long-term (< 15 years) temporal variation in temperate regions. This pattern was consistent across most taxonomic groups, aligning with previous studies that have shown that limited dispersal, environmental heterogeneity, and oceanographic barriers can structure microbial populations at the regional and ocean-basin scales^104,105^.

Seasonal population structuring was evident across most MAGs, with lower F_ST_ values among samples from similar seasons and greater differentiation between those collected in opposite seasons. This indicates cyclical dynamics in population structure, likely driven by fluctuations in temperature, light, or nutrient availability. Interestingly, in all tested MAGs, a modest increase in divergence over time was also observed. This divergence may arise through drift, strain replacement, or local adaptive processes^19,106^. Recently, Hoetzinger and colleagues^88^ reported temporal and spatial increases in F_ST_ for four microbial taxa across five continental waterbodies. They observed changes in F_ST_ ranging from 0.0065 to 0.37 per 1,000 days and from 0.039 to 0.053 per 1,000 km, highlighting measurable divergence over ecological timescales. In comparison, the 389 Mediterranean MAGs analyzed in our study showed more modest F_ST_ values, ranging from 0.001 to 0.09 per 1,000 days, and 0.002 to 0.04 per every 1,000 KM, suggesting a slower rate of population differentiation in coastal marine environments.

The most rapid temporal divergence reported by Hoetzinger *et al.,* was in *Polynucleobacter paneuropaeus*, with a ΔF_ST_ ≈ 0.37 per 1,000 days^88^. In contrast, the highest temporal F_ST_ values in our MAG collection were observed in the family Sphingomonadaceae at BBMO (∼0.07), and in a Flavobacteria UA16 MAG at SOLA (∼0.09), per 1,000 days. These differences suggest that microbial populations in lakes may experience faster genomic divergence than those in coastal marine environments, possibly due to greater isolation of populations.

Nevertheless, both environments contain taxa with minimal divergence. For example, Hoetzinger and colleagues reported that *Candidatus Fonsibacter* (LD12) exhibited a relatively slow F_ST_ increase (∼0.017 per 1,000 days)^88^. Similarly, in our dataset, the marine counterpart to LD12, *Pelagibacteraceae* (SAR11, 9 MAGs), showed low rates of temporal differentiation (∼0.005 in BBMO and ∼0.01 in SOLA over 1,000 days), reinforcing the idea that some cosmopolitan clades maintain high genomic cohesion across time and space. Likewise, long-term observations in Lake Mendota (Madison, USA) also support this trend, revealing gradual shifts in strain composition over two decades^107^.

Altogether, these findings emphasize that microbial population differentiation can unfold over ecologically relevant timescales, with both rapid and slow divergence patterns coexisting depending on lineage and environment. This underlines the value of tracking such long-term dynamics to understand microbial evolutionary responses to ongoing global change^108^.

In conclusion, our population genomics survey of 1,505 prokaryotic MAGs across broad spatiotemporal scales, spanning up to 15 years in the Mediterranean Sea and extending across the global ocean, revealed three main patterns of genomic differentiation: (i) strong differentiation in both space and time, (ii) weak differentiation in both dimensions, and (iii) weak temporal yet strong spatial structure, the last being the most widespread across most taxonomic groups (68.4%). Thus, our study suggests that global-scale geography outweighs long-term seasonality in shaping population differentiation in marine prokaryotes, yet the extent of differentiation remains taxon-specific. As the studied MAGs derive from the Northwestern Mediterranean Sea, extending this framework to genomes from diverse regions and environments will be crucial for testing the generality of these patterns. Altogether, by integrating long-term time series with extensive global surface-ocean metagenomes, we provide a framework for tracking microbial population responses to environmental heterogeneity across large spatiotemporal scales. Collectively, these findings enhance our ability to understand and anticipate how the ocean microbiome will respond to accelerating global change.

## DATA AVAILABILITY

The metagenomic datasets used in this work correspond to publicly available and newly released data. Raw metagenomic sequences for the SOLA time series (Banyuls Bay Microbial Observatory) are available in the European Nucleotide Archive (ENA) under accession numbers PRJEB66489 and PRJEB26919^85^. Metagenomic sequences from the global ocean dataset of the *Tara Oceans* expedition are also available in ENA under project PRJEB402 and PRJEB9740^48,104,109^. New data released in this work includes the metagenomic sequences from the BBMO time series, which is available in ENA under project PRJEB51979, and the BBMO Prokaryotic Metagenome Assembled Genome catalog v1, available in Zenodo under DOI: 10.5281/zenodo.17159964. Additional data and results produced in this work, such as read alignment files (CRAM), MAG abundances and F_ST_ tables, and all supplementary graphs produced for each genome are available in Zenodo under DOI: 10.5281/zenodo.17634504.

## ACKNOWLEDGMENTS

We thank all members of the LTER BBMO (http://bbmo.icm.csic.es/) and SOLA time-series teams for their sustained efforts and contributions over the years to maintaining both observatories, as well as the *Tara Oceans* Consortium for providing publicly available oceanomics data. Bioinformatics analyses were performed at the MARBITS platform of the Institut de Ciències del Mar (ICM) [https://marbits.icm.csic.es] and Centro de Supercomputación de Galicia (CESGA; https://www.cesga.es/). FL was supported by the Spanish National Program FPI 2016 (BES-2016-076317, MICINN, Spain). RL was supported by a Ramón y Cajal fellowship (RYC-2013-12554, MINECO, Spain). RM was supported by EPIC (PID2022-137508NB-I00, MICINN). This work was supported by the project INTERACTOMICS (CTM2015-69936-P, MINECO, Spain), MicroEcoSystems (240904, RCN, Norway), MINIME (PID2019-105775RB-I00, MICINN, Spain), and MAORI (PID2022-136281NB-I00, MICIU) to RL. This research was co-funded by the European Union (GA#101059915 - BIOcean5D). Views and opinions expressed are however those of the author(s) only and do not necessarily reflect those of the European Union. Neither the European Union nor the granting authority can be held responsible for them.

## SUPPLEMENTARY MATERIAL

**Figure S1.**
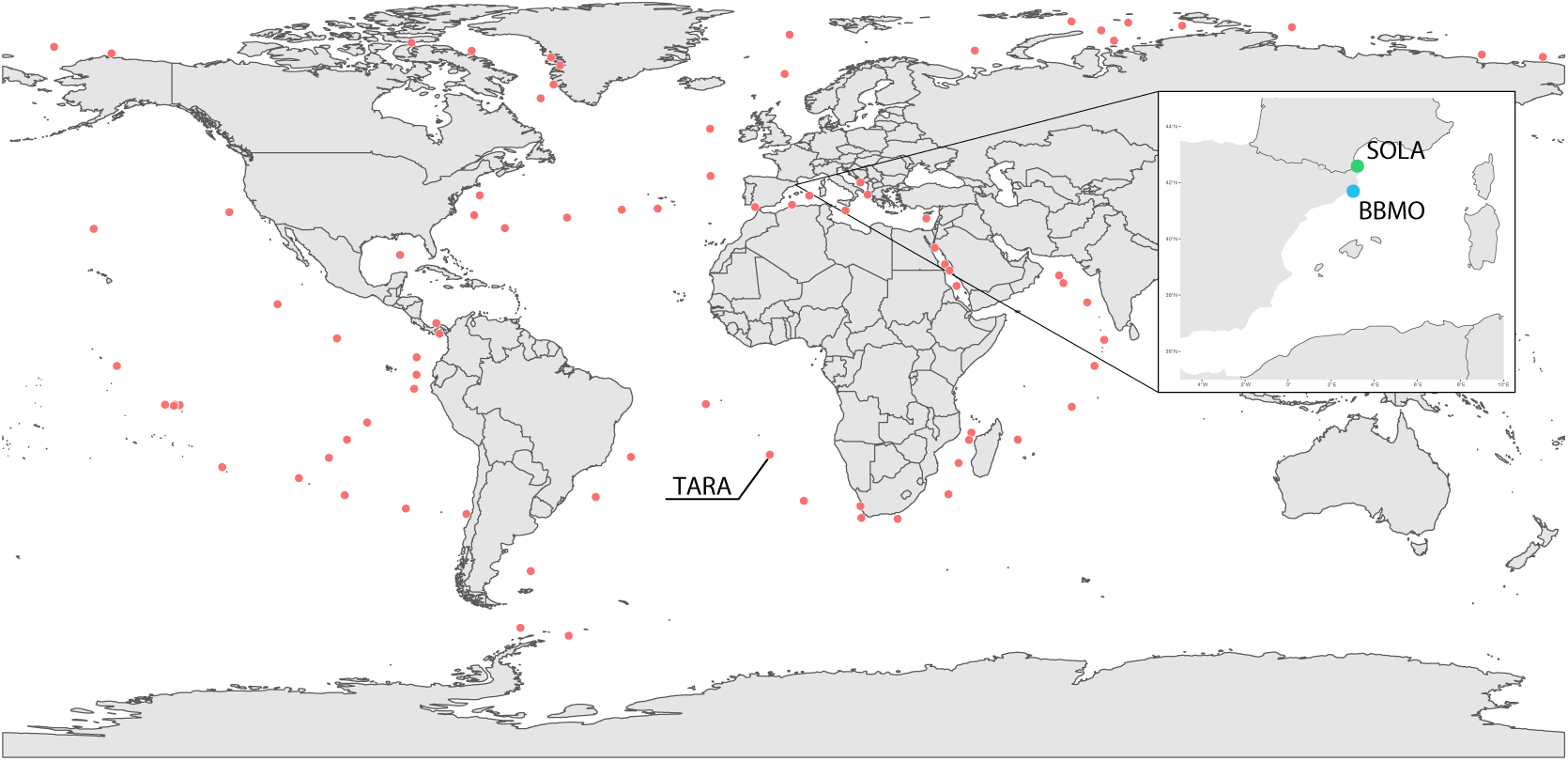
Locations of all Samples from BBMO (blue), SOLA (green), and TARA (red).

**Figure S2.**
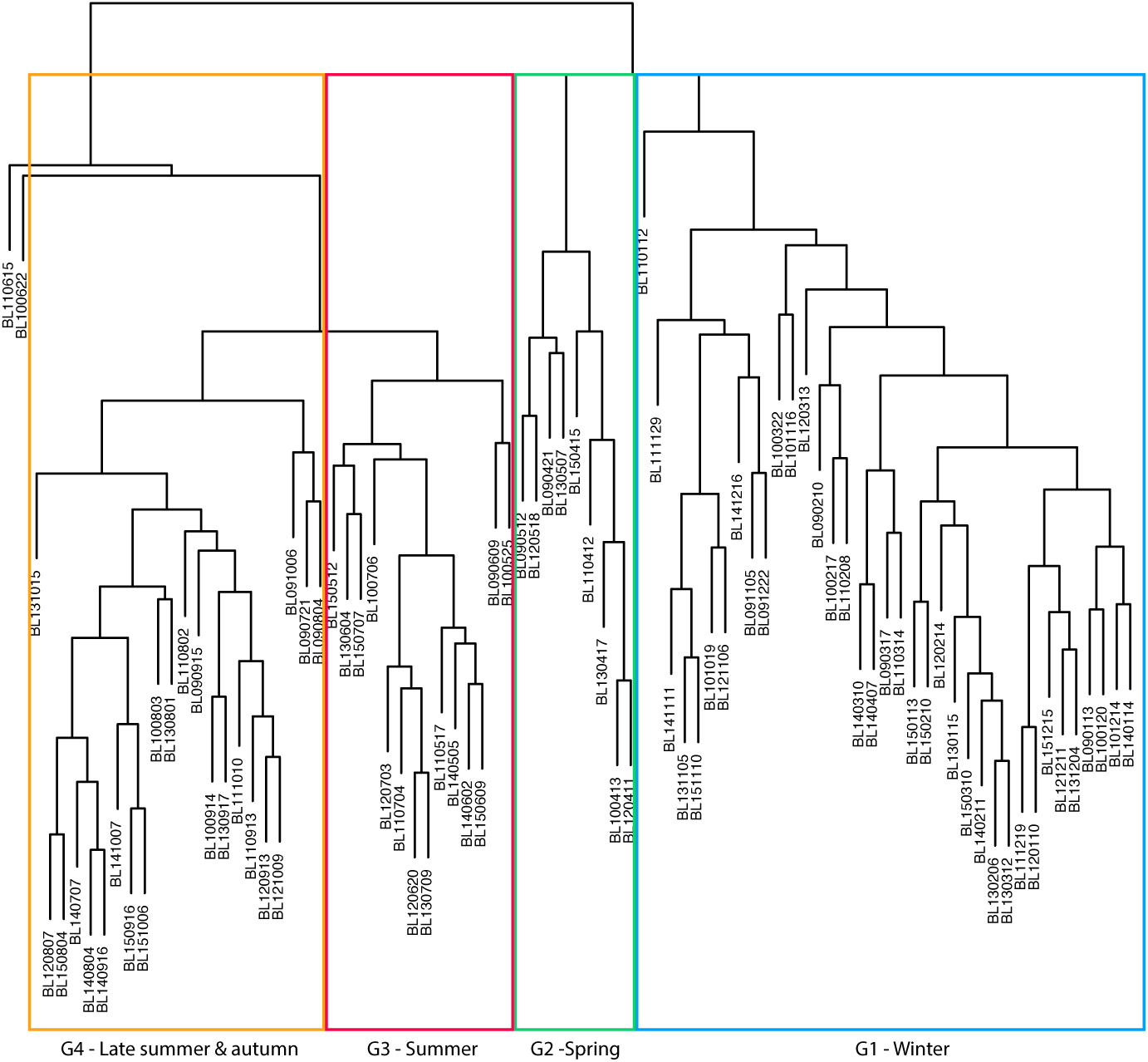
UPGMA clustering of Bray-Curtis dissimilarities among the 84 BBMO metagenomes calculated using SIMKA (kmer = 21).

**Figure S3.**
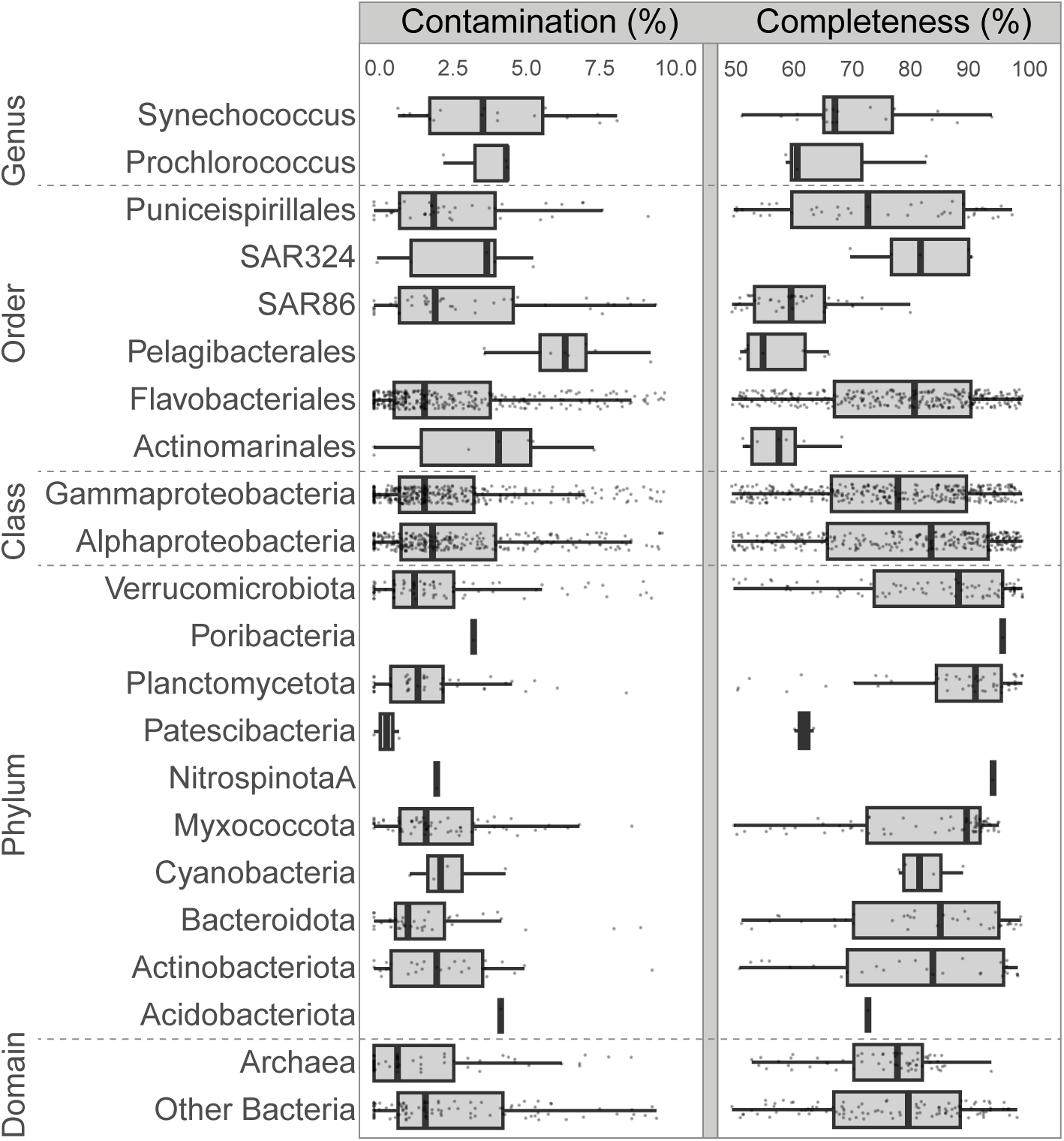
Boxplots showing the completeness and contamination of the 1,505 MAGs, grouped by different taxonomic affiliations. Taxonomic levels range from Genus to Domain, highlighting specific ecologically relevant groups. MAGs belonging to multiple levels were assigned only to the lowest rank (*e.g., Prochlorococcus* MAGs were not included in the Cyanobacteria or Bacteria categories).

**Figure S4.**
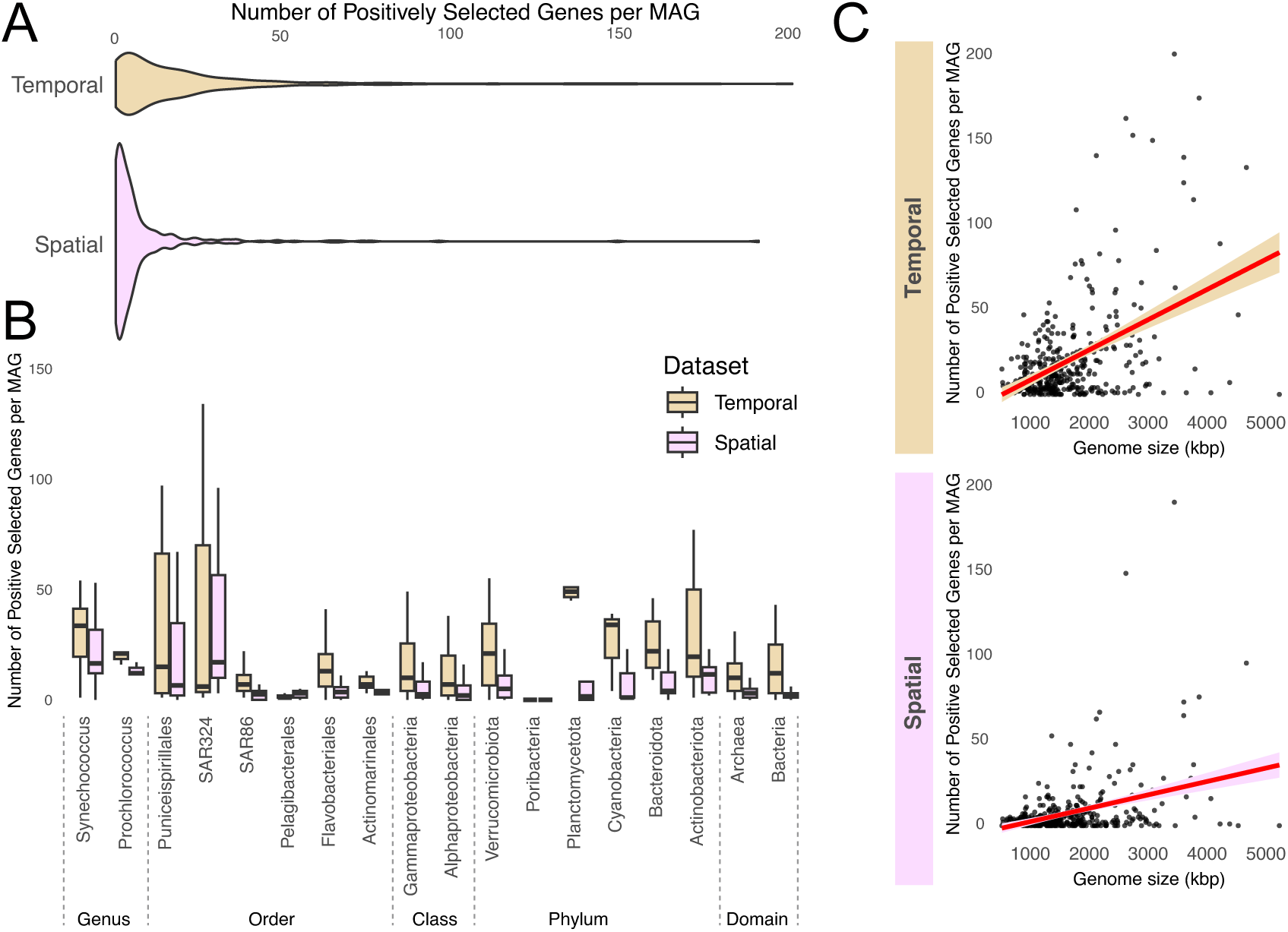
Number of positively selected genes per MAG across (A) datasets and (B) taxonomic groups. Taxonomic levels range from Genus to Domain, highlighting specific ecologically relevant groups. **(C)** Number of positively selected genes per MAG in relation to genome size (kbp) for temporal (BBMO and SOLA) and spatial (TARA Oceans) datasets. Each dot represents one MAG. Results are presented separately for spatial and temporal datasets to facilitate the comparison of patterns across different scales.

**Figure S5.**
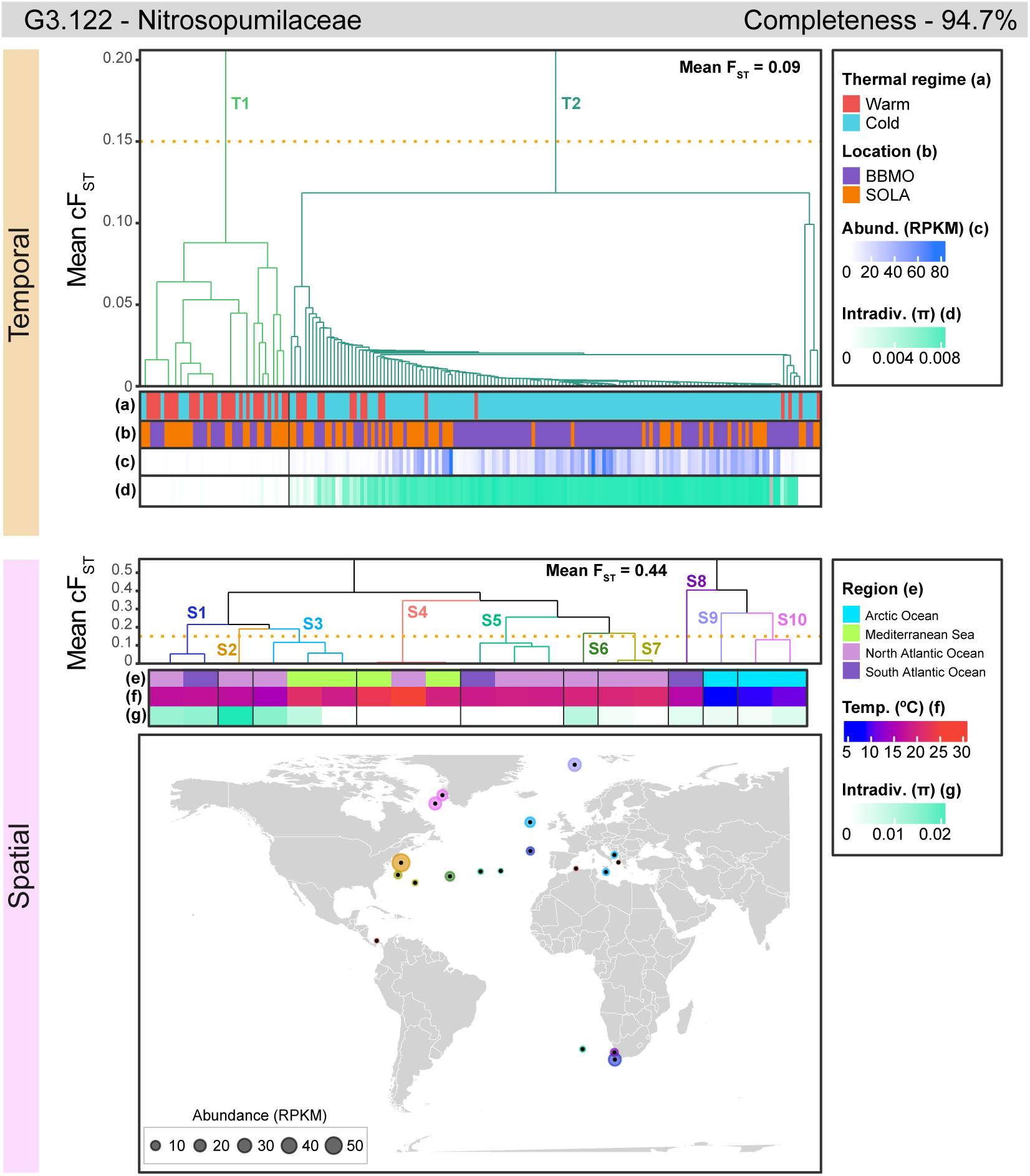
Population structure in the two time series (BBMO and SOLA) and the global ocean (TARA) for Nitrosopumilaceae MAG G3.122. Potential populations were defined using a Mean cF_ST_ threshold of 0.15, computed with the UPGMA clustering algorithm (dendrogram axis), which represents the average F_ST_ for each cluster. In turn, the Mean F_ST_ value indicates the overall mean of all pairwise F_ST_ values for each genome within each dataset. Different colors and labels distinguish between potential populations, labeled as T for Temporal and S for Spatial datasets. For each dataset, the main thermal regimes (Warm vs. Cold waters), location (BBMO vs. SOLA), abundance (RPKM, reads per kilobase of genome and million mapped reads), intradiversity (π), and temperature (°C) are displayed. Missing values are shown in grey. Genome completeness is also reported.

**Figure S6.**
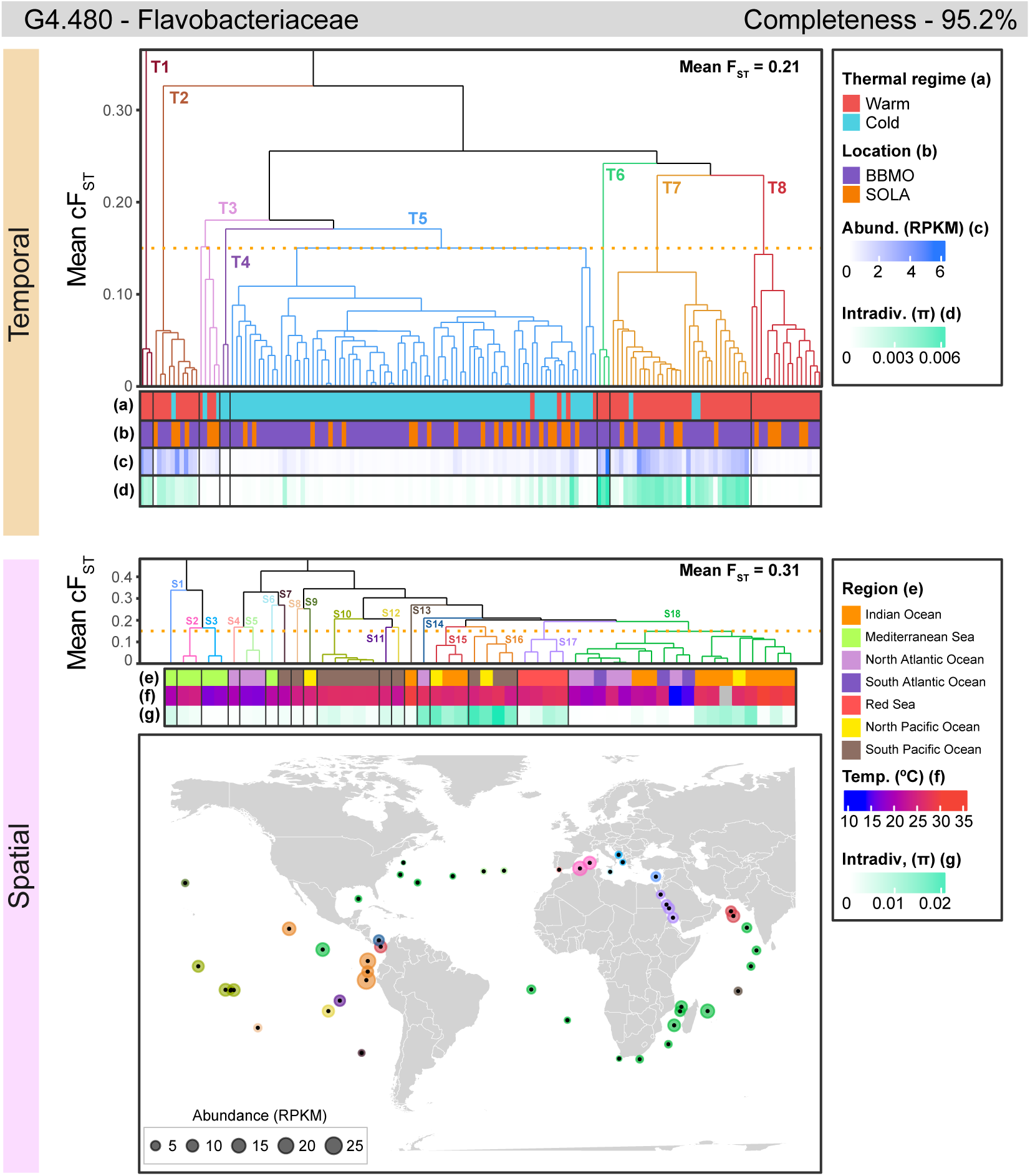
Population structure in the two time series (BBMO and SOLA) and the global ocean (TARA) for Flavobacteriaceae MAG G4.480. Potential populations were defined using a Mean cF_ST_ threshold of 0.15, computed with the UPGMA clustering algorithm (dendrogram axis), which represents the average F_ST_ for each cluster. In turn, the Mean F_ST_ value indicates the overall mean of all pairwise F_ST_ values for each genome within each dataset. Different colors and labels distinguish between potential populations, labeled as T for Temporal and S for Spatial datasets. For each dataset, the main thermal regimes (Warm vs. Cold waters), location (BBMO vs. SOLA), abundance (RPKM, reads per kilobase of genome and million mapped reads), intradiversity (π), and temperature (°C) are displayed. Missing values are shown in grey. Genome completeness is also reported.

